# Two pathways for thiosulfate oxidation in the alphaproteobacterial chemolithotroph *Paracoccus thiocyanatus* SST

**DOI:** 10.1101/683490

**Authors:** Moidu Jameela Rameez, Prosenjit Pyne, Subhrangshu Mandal, Sumit Chatterjee, Masrure Alam, Sabyasachi Bhattacharya, Nibendu Mondal, Jagannath Sarkar, Wriddhiman Ghosh

## Abstract

Chemolithotrophic bacteria oxidize various sulfur species for energy and electrons, thereby operationalizing biogeochemical sulfur cycles in nature. The best-studied pathway of bacterial sulfur-chemolithotrophy, involving direct oxidation of thiosulfate to sulfate (without any free intermediate) by the SoxXAYZBCD multienzyme system, is apparently the exclusive mechanism of thiosulfate oxidation in facultatively chemolithotrophic alphaproteobacteria. Here we explore the molecular mechanisms of sulfur oxidation in the thiosulfate- and tetrathionate-oxidizing alphaproteobacterium *Paracoccus thiocyanatus* SST, and compare them with the prototypical Sox process characterized in *Paracoccus pantotrophus*. Our results revealed the unique case where, an alphaproteobacterium has Sox as its secondary pathway of thiosulfate oxidation, converting ∼10% of the thiosulfate supplied whilst 90% of the substrate is oxidized via a Tetrathionate-Intermediate pathway. Knock-out mutation, followed by the study of sulfur oxidation kinetics, showed that thiosulfate-to-tetrathionate conversion, in SST, is catalyzed by a thiosulfate dehydrogenase (TsdA) homolog that has far-higher substrate-affinity than the Sox system of this bacterium, which, remarkably, is also less efficient than the *P. pantotrophus* Sox. *soxB*-deletion in SST abolished sulfate-formation from thiosulfate/tetrathionate while thiosulfate-to-tetrathionate conversion remained unperturbed. Physiological studies revealed the involvement of glutathione in SST tetrathionate oxidation. However, zero impact of the knock-out of a thiol dehydrotransferase (*thdT*) homolog, together with no production of sulfite as an intermediate, indicated that tetrathionate oxidation in SST is mechanistically novel, and distinct from its betaproteobacterial counterpart mediated by glutathione, ThdT, SoxBCD and sulfite:acceptor oxidoreductase. All the present findings collectively highlight extensive functional diversification of sulfur-oxidizing enzymes across phylogenetically close, as well as distant, bacteria.

## 1. Introduction

Chemolithotrophic sulfur oxidation is a primordial metabolic process central to our understanding of early life on Earth (Ghosh and Dam, 2009). In nature, sulfur exists in a wide range of oxidation states (−2 to +6), and in doing so passes through a complex variety of biogeochemical processes (Luther et al., 1985). Sulfur cycling, in diverse ecosystem, essentially involves sulfur compounds syntrophy (i.e., inter-microbial transfer of various redox states of sulfur), physicochemically or biochemically intermingled with the transformation dynamics of carbon, nitrogen, iron and other metals (Cutter and Kluckhohn 1999; Mopper and Kieber 2002; Owens et al. 2016). Among the various reduced states of sulfur, thiosulfate constitutes a key junction in the network of sulfur-species transformations in diverse ecosystems, such as marine sediments (Jorgensen, 1990a; Thamdrup et al., 1994), freshwater sediments (Jorgensen, 1990b), meromictic lakes (Kondo et al., 2000) and hot spring waters (Xu et al., 1998). In these biogeochemically diverse environments, taxonomically diverse microorganisms oxidize/reduce thiosulfate to meet their bioenergetic requirements; among them, the thiosulfate-oxidizing chemolithotrophs are the major players of the oxidative half ot the sulfur cycle.

Distinct sets of enzymes and electron transport systems are known to render the oxidation of thiosulfate in phylogenetically diverse bacteria and archaea (Ghosh and Dam, 2009). Of the various pathways of thiosulfate oxidation, the SoxXAYZBCD multienzyme system-mediated mechanism epitomized in chemo-/photo-lithotrophic alphaproteobacteria (Kelly et al., 1997; Friedrich et al., 2000; Mukhopadhyaya et al., 2000; Appia-Ayme et al., 2001; Friedrich et al., 2001, 2005; Kappler et al., 2001; 2004; Bamford et al., 2002; Sauve et al., 2007; Ghosh and Dam, 2009) is one of the best studied. Here, the sulfane sulfur atom of thiosulfate is first oxidatively coupled with the SoxY-cysteine-sulfhydryl group of the SoxYZ complex by the action of the cytochromes SoxXA, thereby forming a SoxZY-cysteine S-thiosulfonate adduct; subsequently, the sulfone as well as sulfane sulfurs of the SoxYZ bound thiosulfate is oxidized to sulfate by the sequential activity of SoxB, Sox(CD)_2_, and then again SoxB (Kelly et al., 1997; Appia-Ayme et al., 2001; Friedrich et al., 2001; Quentmeier and Friedrich, 2001; Bamford et al., 2002). In this context it is noteworthy that despite *sox* structural genes being widespread in Bacteria, the typical Sox mechanism of direct 8-electron transfer (to the electron transport systems) during the oxidation of thiosulfate (without the formation of any free intermediate) has thus far been established only in the alphaproteobacterial lithotrophs, which apparently have no other pathway for thiosulfate oxidation. Only a small number of chemolithotrophic magnetotactic alphaproteobacteria are known to utilize a truncated SoxXAYZB system to oxidize thiosulfate in the same way as the anaerobic and anoxygenic photolithotrophs do (i.e. in collaboration with reverse-acting sulfate-reducing enzymes) (Eisen et al., 2002; Verte et al., 2002; Dahl, 2008; Grimm et al., 2008).

In contrast to the alphaproteobacteria, thiosulfate oxidation in most of the beta- and gammaproteobacterial chemolithotrophs proceeds via the formation of tetrathionate-intermediate (S_4_I) (Visser et al., 1996; Bugaytsova and Lindström, 2004; Ghosh et al., 2005; Dam et al., 2007; Rzhepishevska et al., 2007; Kikumoto et al., 2013; Pyne et al., 2017, 2018). In these bacteria, thiosulfate to tetrathionate conversion is mediated either by (i) *doxDA*-encoded thiosulfate:quinone oxidoreductase (Rzhepishevska et al., 2007; Kikumoto et al., 2013) that is also present in thermoacidophilic archaea (Müller et al., 2004;), or by (ii) thiosulfate dehydrogenase (Pyne et al., 2018) that is also present in bacteria incapable of further oxidation of tetrathionate to sulfate (Hensen et al., 2006; Denkmann et al., 2012; Frolov et al., 2013; Brito et al., 2015; Orlova et al., 2016). The S_4_I formed in this way is subsequently oxidized either (i) by the activity of the pyrroloquinoline quinone (PQQ)-binding tetrathionate hydrolase (TetH), as described in *Acidithiobacillus* species (De Jong et al.,1997a; Kanao et al., 2007; Rzhepishevska et al., 2007; van Zyl et al., 2008; Kanao et al., 2013), or (ii) via coupling with glutathione GSH (to form the glutathione:sulfodisulfane adduct GS–S–S–SO_3_^−^ and sulfite) by the action of another PQQ-binding protein called thiol dehydrotransferase (ThdT, which is a homolog of the XoxF variant of methanol dehydrogenase), followed by the oxidation of GS–S–S–SO_3_^−^ via iterative typical actions of SoxB and SoxCD, as reported in *A. kashmirensis* (Pyne et al., 2018).

The current state of knowledge recognizes the preminence/monopoly of the Sox pathway in alphaproteobacterial oxidation of thiosulfate to sulfate, while the S_4_I is essentially confined to the beta and gammaproteobacteria [the S_4_I pathway has, thus far, been implicated only in one alphaproteobacterium, *Acidiphilium acidophilum* (Meulenberg et al., 1993; De Jong et al., 1997b), and that is the only potential presence of this pathway outside Beta- and Gammaproteobacteria]. Counter to this existing scenario, the present molecular investigation of the chemolithotrophic machinery of *Paracoccus thiocyanatus* strain SST (= LMG 22699 = MTCC 7821; see Ghosh and Roy, 2007) reveals a unique case where, an alphaproteobacterium has Sox as its secondary pathway of thiosulfate oxidation while the S_4_I pathway is the frontline mechanism for this purpose. Functional dynamics and intersections of the Sox and S_4_I pathways of SST were delineated via physiological studies, whole genome sequencing and analysis, and recombination-based knock-out mutagenesis of putative sulfur oxidation genes. Results were then considered in the context of the prototypical Sox and S_4_I pathways of *Paracoccus pantotrophus* (Friedrich et al., 2001; Quentmeier and Friedrich, 2001; Sauve et al., 2007) and *Advenella kashmirensis* (Ghosh et al., 2005; Dam et al., 2007; Pyne et al., 2017, 2018) repsectively.

## 2. Materials and methods

### 2.1. Bacterial strains, media, and culture conditions

All strains, whether wild-type or engineered (see Table 1), were routinely grown or maintained in Luria Bertani (LB) medium, which was supplemented with appropriate antibiotics as and when required. Chemolithoautotrophic properties of *P. thiocyanatus* SST wild-type, or its mutants, were tested in modified basal and mineral salts (MS) solution (1 g NH_4_Cl, 4 g K_2_HPO_4_, 1.5 g KH_2_PO_4_, 0.5 g MgCl_2_, and 5 mL trace metals solutionL^-1^ distilled water) supplemented with 20 mM sodium thiosulfate (MST) or 10 mM potassium tetrathionate (MSTr); in this way, both MST (pH 7.0) and MSTr (pH 7.0/ 8.0 as per experimental requirement) contained sulfur equivalent to 40 mM S (Ghosh et al., 2005). For chemoorganotrophic growth, MS was supplemented with 1.0 g L^-1^dextrose (MSD, pH 8.0). *P. thiocyanatus* strains were grown at 28°C, whereas *Escherichia coli* strains used in genetic experiments were grown at 37°C. Antibiotics were used at the following concentrations: ampicillin, 100 µg mL^−1^; kanamycin, 40 µg mL^−1^; streptomycin 100 µg mL^−1^; and rifampicin 50 µg mL^−1^.

**Table 1.** . Bacterial strains and plasmids used in this study.

| Strain | Genotype / characteristic(s) | Reference /source |
| --- | --- | --- |
| *Escherichia coli* S17-1 λpir | Thi-Pro-Hsd-recA-lacZ::RP4-2 (tet::Mu. kan:: Tn7 [Tp<sup>R</sup>, S1-mR]), λpir | Simon et al., 1983 |
| *Paracoccus thiocyanatus* SST | Wild-type , Str<sup>R</sup> | Ghosh and Roy, 2007 |
| *Paracoccus thiocyanatus* SSTR | Spontaneous Rif<sup>R</sup> mutant of *P. thiocyanatus* SST; Str<sup>R</sup>, Rif<sup>R</sup> | In this study |
| Pt_KoT | Δ*tsdA* knock-out mutant of *P. thiocyanatus* SSTR; Kan<sup>R</sup>, Sm<sup>R</sup>, and Rif<sup>R</sup> | In this study |
| Pt_KoB | Δ*soxB* knock-out mutant of *P. thiocyanatus* SSTR; Str<sup>R</sup>, Rif<sup>R</sup>, Kan<sup>R</sup> | In this study |
| Pt_KoX | *xoxF::Kan<sup>R</sup>* knock-out mutant of *P. thiocyanatus* SSTR; Str<sup>R</sup>, Rif<sup>R</sup>, Kan<sup>R</sup> | In this study |
| **Plasmid / construct** | **Genotype** | **Reference /source** |
| pKAS32 | Amp<sup>R</sup>, *rpsL*, R6K, RP4 | Skorupski and Taylor, 1996 |
| pSUP5011::Tn5-*mob* | pBR325(BamHI)::Tn5-*mob*, Amp<sup>R</sup>, Kan<sup>R</sup>, (Cam<sup>R</sup>) | Simon et al., 1983 |
| pKKoT | pKAS32::KoT; Amp<sup>R</sup> | This study |
| pKKoT::Kan<sup>R</sup> | KoT::Kan<sup>R</sup> cloned in PKAS32; Amp<sup>R</sup>, Kan<sup>R</sup> | This study |
| pKKoB | pKAS32::KoB; Amp<sup>R</sup> | This study |
| pKKoB::Kan<sup>R</sup> | KoB::Kan<sup>R</sup> cloned in PKAS32; Amp<sup>R</sup>, Kan<sup>R</sup> | This study |
| pKKoX | pKAS32::KoX; Amp<sup>R</sup> | This study |
| pKKoX::Kan<sup>R</sup> | KoX::Kan<sup>R</sup> cloned in PKAS32; Amp<sup>R</sup>, Kan<sup>R</sup> | This study |

### 2.2. Physiological / biochemical experiments and analytical methods

For all batch culture experiments in MS-based media, 100 mL test media were inoculated with 1% (v/v) LB-cultures having OD_600_ of 0.6. Samples were collected from the test cultures at appropriate time points, and cells were removed from them by centrifugation before pH of the spent media, and concentrations of thiosulfate/tetrathionate/sulfite/sulfate, were recorded. Concentrations of dissolved thiosulfate, tetrathionate and sulfate in spent media were measured by iodometric titration, cyanolytic method and gravimetric precipitation method respectively (Kelly et al., 1994; Alam et al., 2013). Sulfite was measured spectrophotometrically with pararosaniline as the indicator (West and Gaeke, 1956). To corroborate the results obtained by the above methods, sulfate, sulfite and thiosulfate were additionally quantified by anion chromatography via chemical suppression, using an Eco IC (Metrohm AG, Switzerland) equipped with a conductivity detector (Metrohm, IC detector 1.850.9010) and autosampler (863 Compact Autosampler, Switzerland).

Separation of ions was carried using a Metrosep A Supp5 - 250/4.0 (6.1006.530) anion exchange column (Metrohm AG); a mixed solution of 1.0 mM sodium hydrogen carbonate and 3.2 mM sodium carbonate was used as the eluent; 100 mM sulfuric acid was used as the regenerant; flow rate was 0.7 mL min ^-1^, and injection volume 100 µL. Samples were diluted 1000-fold with deionized water (Siemens, <0.06 μS), and filtered by passing through 0.22 µm hydrophilic polyvinylidene fluoride membranes (Merck Life Science Private Limited, India), prior to analysis. Sigma-Aldrich (USA) standard chemicals were used to prepare the calibration curve for quantification. Overall sample reproducibility was ± 0.2 ppm.

Sulfur-substrate-dependent O_2_ consumption rate was measured in SST cells grown in MST (pH 7.0), MSTr (pH 8.0) or MSD (pH 8.0) medium. MST-grown cells were harvested when 80% of the supplied thiosulfate had been converted to tetrathionate, while MSTr-grown cells were harvested when 50% of the supplied tetrathionate had been converted to sulfate. MSD-grown cells were harvested when the culture was in mid-log phase of growth. Cells were harvested, washed twice with, and resuspended in, potassium phosphate buffer (50 mM, pH 8.0). Sulfur-compounds-dependent O_2_ consumption rates of whole cells were determined under varying pH conditions (4.5 to 10.5), using a biological O_2_ monitor having a Clark-type O_2_ electrode (Yellow Springs Instruments, Ohio, USA). The 3.0 mL assay mixture contained 50 mM of the assay buffer specific for the pH at which oxidation was measured (see below), washed-cell suspension (equivalent to 100 µg of whole cell protein), and sodium thiosulfate (10 mM) / potassium tetrathionate (5 mM) / sodium sulfite (2 mM) in 5 mM EDTA. Reactions were started by adding the sulfur substrate. O_2_ consumption rates, expressed as nmol O_2_ consumed mg cellular protein^-1^ min^-1^, were corrected for chemical reactions and endogenous respiration. The different assay buffers used for the different pH ranges were as follows: sodium acetate–acetic acid buffer for pH 4.5-6.0 (Gomori, 1955), potassium phosphate buffer for pH 6.0-8.5 (Sambrook and Russell, 2001) and glycine–sodium hydroxide buffer for pH 8.5-10.5 (Gomori, 1955). Specific pH values of the individual assay mixtures were set by mixing the components of the respective buffers in specific ratios described in the corresponding literature.

Effects of the thiol-inhibitor N-ethyl maleimide (NEM) and the thiol-group-containing tripeptide glutathione (GSH) on the substrate-dependent O_2_ consumption rates in the presence of sodium thiosulfate (10 mM) / potassium tetrathionate (5 mM) / sodium sulfite (2 mM) were measured essentially as above: specifically, the 50 mM assay buffer had pH 8.5, and the washed cells were incubated in 1 mM NEM or 10 mM GSH for 10 minutes before the assay. All calculations pertaining to O_2_ consumption rates were done based on the assumption (according to the instrument manual) that an 18% change in O_2_ concentration equals to 2.71 µl of O_2_ consumed.

### 2.3. Formulation of a pH dependence factor

A pH-dependence factor (*F*_*pH*_), based on the sulfur-substrate-dependent O_2_ consumption rates (*R*) recorded at various pH values (4.5 - 10.5), was formulated to quantitatively compare the extent to which the thiosulfate-and tetrathionate-oxidizing machineries of *P. thiocyanatus* SST were influenced by changes in the pH of the chemical milieu. The entire pH range (4.5 to 10.5) over which O_2_ consumption was recorded in the presence of thiosulfate/tetrathionate was divided into 12 classes (4.5-5.0, 5.0-5.5, 5.5-6.0, through 10.0-10.5), each of which encompassed a fixed pH window of 0.5. Change in the O_2_ consumption rate (Δ*R*) for a particular substrate, across any pH window of 0.5, was first calculated by dividing the *R* value obtained at the upper pH-limit of the window with the *R* value obtained at the lower pH-limit of the window (equation 1). Following this, Δ*R* – 1 gave the pH dependence factor (*F*_*pH*_) of the oxidation process, over the pH window considered (equation 2). An *F*_*pH*_ value becomes 0 when the efficacy of the process is not influenced by pH change within the window under consideration; in contrast, *F*_*pH*_ values > 0 denote increased efficiency with increasing pH, and *F*_*pH*_ values < 0 denote impairment of the process wth increasing pH, within the window considered. For a particular process, collation of the *F*_*pH*_ values obtained for all contiguous pH windows gave the extent of its pH-dependence.

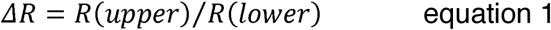

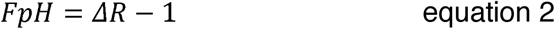

### 2.4. Whole genome sequencing and annotation

Whole genome sequencing for *P. thiocyanatus* SST was carried out on genomic DNA isolated form stationary phase cultures of this bacterium. Two separate sequencing runs on the Ion Torrent Personal Genome Machine (Ion PGM) and the Ion Proton platform (both from Thermo Fisher Scientific, Waltham, MA, USA) were conducted for this purpose with 400 bp and 200 bp read chemistry respectively. First, the 1,054,074 high-quality reads obtained from the Ion PGM run (mean read length 258 bp) were used in assembly using Newbler 2.8 (454 Sequencing System Software, Roche; http://www.454.com). This yielded 510 contigs, which constituted a consensus of 3,465,907 bp. These were subsequently used as trusted contigs in a co-assembly [using SPAdes 3.11.1 (Bankevich et al., 2012)] with the Ion PGM reads, plus the 4,675,076 high-quality reads (mean read length 163 bp) obtained from the Ion Proton run. This co-assembly resulted in the reduction of contig count to 289 with a total consensus of 3,530,182 bp. Of these, 282 contigs that were > 200 bp long were annotated using the automated Prokaryotic Genome Annotation Pipeline (PGAP) of the National Center for Biotechnology Information (NCBI; Bethesda, MD, USA) and deposited at DDBJ/ENA/GenBankunder the accession QFCQ00000000, the version described in this paper is QFCQ01000000. All information regarding this Whole Genome Shotgun project is available in the GenBank under the BioProject PRJNA464267, while the raw sequence reads obtained from the Ion PGM and Ion Proton runs are available in the NCBI Short Read Archive (SRA) under the run accession numbers SRX4046611 and SRX4046610 respectively.

Completeness of the *P. thiocyanatus* SST genome was determined using CheckM 1.0.12 (Parks et al., 2015). For this purpose, a custom database of phylogenetic marker genes specific for the genus *Paracoccus* was first constructed from the set of marker genes provided with CheckM. Parallel to this, open reading frames (ORFs) were predicted wthin the SST genome using the software Prodigal, and then homologs of marker genes encompassed in the custom database were searched in the Prodigal-derived ORF/gene catalog using HMMER algorithm. Completeness of the SST genome was calculated based on the number of *Paracoccus*-specific markers detected in the gene-catalog.

Homologs of genes involved in bacterial sulfur oxidation were searched either manually in the PGAP-annotated SST genome or by mapping/aligning query gene sequences from other alphaproteobacterial species onto the *P. thiocyantus* SST genome sequence using the PROmer utility of the MUMmer 3.23 package (Kurtz et al., 2004). Annotations of the ORFs identified by the latter method were corroborated by NCBI BlastP analysis of the putative amino acid sequences, followed by their conserved domain search with reference to the Conserved Domain Database (CDD).

### 2.5. Knock-out mutagenesis

Homologous recombination-based techniques were used to render deletion mutations in the *tsdA* (locus tag DIE28_04655) and *soxB* (locus tag DIE28_12735) genes, and insertional inactivation of the *xoxF* gene (locus tag DIE28_01705) of SST. To knock-out *tsdA*, the upstream and downstream flanking regions of the ORF were amplified by two separate PCR reactions using the primer-pairs (i) Pt_Ko_tsdA_F and Pt_Ko_tsdA_FU_R, and (ii) Pt_Ko_tsdA_FU_F and Pt_Ko_tsdA_R (sequences given in Supplementary Table S1). The PCR products obtained (see the steps denoted as PCR1 and PCR2 in Supplementary Fig. S1) were used as templates for overlap extension PCR with the primer set Pt_Ko_tsdA_F and Pt_Ko_tsdA_R to generate a 1944 bp fused product (KoT) which encompassed both the flanking regions (see the step denoted as PCR3 in Supplementary Fig. S1). This product was then cloned into pKAS32 between the *Xba*I and *Eco*RI sites to generate the construct pKKoT. Simultaneously, the *kanR* gene with its upstream promoter region (i.e. the Kan^R^ marker cartridge) was PCR amplified from the plasmid pSUP5011::Tn5-mob, without any terminator sequence, using the primer-pair kanR_KpnI_F and kanR_KpnI_R (sequences given in Supplementary Table S1). The Kan^R^ cartridge was then inserted within the *Kpn* I restriction site of pKKoT, which in turn was introduced through PCR using the primer-pair Pt_Ko_tsdA_FU_F and Pt_Ko_tsdA_R (see the step denoted as PCR2 in Supplementary Fig. S1). This gave rise to the final construct pKKoT::Kan^R^.

To knock-out *soxB*, the upstream and downstream flanking regions of theORF were amplified by two separate PCR reactions using the primer-pairs (i) Pt_Ko_soxB_F and Pt_Ko_ soxB _FU_R, and (ii) Pt_Ko_soxB_FU_F and Pt_Ko_soxB_R (sequences given in Supplementary Table S2). The PCR products obtained (see the steps denoted as PCR1 and PCR2 in Supplementary Fig. S2) were used as templates for overlap extension PCR with the primer set Pt_Ko_soxB_F and Pt_Ko_soxB_R to generate a 2345 bp fused product (KoB) which encompassed both the flanking regions (see the step denoted as PCR3 in supplementary Fig. S2). This product was then cloned into pKAS32 between the *Xba*I and *Sac*I sites to generate the construct pKKoB. The Kan^R^ cartridge, obtained as described abve, was then inserted within the *Kpn* I restriction site of pKKoB, which in turn was introduced through PCR using the primer-pair Pt_Ko_soxB_FU_F and Pt_Ko_soxB_R (see the step denoted as PCR2 in Supplementary Fig. S2). This gave rise to the final construct pKKoB::Kan^R^.

*xoxF* was mutated by inserting the above mentioned Kan^R^ cartridge into the ORF. Two separate PCRs using the primer-pairs (i) Pt_Ko_xoxF_F and Pt_Ko_xoxF_FU_R, and (ii) Pt_Ko_xoxF_FU_F and Pt_Ko_xoxF_R (Supplementary Table S3), yielded two, 561 bp and 621 bp long, fragments having terminal 30 bp regions complementary to each other, and containing a common *Kpn*I restriction site introduced via the primers Pt_Ko_xoxF_FU_R and Pt_Ko_xoxF_FU_F. Overlap extension PCR (this is denoted as PCR3 in Supplementary Fig. S3) using a mixture of the 561 and 621 bp fragments as template and Pt_Ko_xoxF_F and Pt_Ko_xoxF_R as the primers, yielded a 1152 bp region (KoX) of *xoxF* where the *Kpn*I site got inserted inside the ORF, and an *Xba*I and a *Sac*I site got introduced at the two ends. This product was then inserted into pKAS32 between the latter’s *Xba*I and *Sac*I restriction sites, thereby generating the construct pKKoX. Kan^R^ cartridge was then inserted within *Kpn*I restriction site of pKKoX. This gave rise the final construct pKKoX::Kan^R^.

The mobilizable constructs pKKoT::Kan^R^, pKKoB::Kan^R^ and pKKoX::Kan^R^ were transformed into *Escherichia coli* S17-1 λ*pir* and used as donor strains in conjugative plate mating for transferring corresponding gene replacement cassettes to *P. thiocyanatus* SST. A spontaneous mutant strain *P. thiocyanatus* SSTR, which was resistant to Rifampicin (50 µg mL^-1^) was used as the recipient in the conjugation experiments. Cells from mid-log phase cultures of both the donor and the recipient were harvested and washed twice with 10 mM MgSO_4_ and resuspended in the same. Donors and recipients were then mixed at a 1:2 cell mass ratio and immobilized onto a membrane filter with pore size of 0.2 μm. The membrane filter was placed on an LA plate containing 1% agar, and incubated at 30^°^C for 18 h. Cell mass growing on the membrane was harvested and dissolved in 2 mL 0.9% NaCl solution; ths was subsequently spread at 10^-1^ dilution on LA plates containing both rifampicin (50 µg mL^-1^) and kanamycin (40 µg mL^-1^). Recipients which had successfully exchanged their relevant genomic loci with gene replacement cassettes loaded on the constructs were screened for double cross-over mutants by positively selecting for the cartridge-encoded kanamycin resistance trait and loss of streptomycin sensitivity conferred by the *rpsL* gene of pKAS32. Whilst the resultants Δ*tsdA*, Δ*soxB* and *xoxF*::Kan^R^ mutants were designated as Pt_KoT, Pt_KoB and Pt_KoX respectively, orientations of the antibiotic resistance cartridges with respect to the transcriptional directions of the ORFsmutated were checked by PCR-amplification and sequencing of DNA fragments containing specific genome-cartidge intersections (see Supplementary Fig. S1-S3).

## 3. Results and discussion

### 3.1. *P. thiocyanatus* SST oxidizes thiosulfate via S_4_I formation

When *P. thiocyanatus* SST was grown in MST medium (initial pH 7.0), only 5 mM S sulfate was found to have produced after 12 h incubation (Fig. 1A). During this period of incubation, thiosulfate was continuously depleted and tetrathionate went on accumulating in the medium at a stoichiometry slightly less than the amount of thiosulfate disappearing, which was attributable to the formation of sulfate; at the end of 12 h, this difference was equivalent to 5 mM S, and the same was detected fully in the form of sulfate. In this way, at the 12^th^ hour, no thiosulfate remained in the medium, and 35 mM S tetrathionate was there in addition to the 5 mM S sulfate, with concomitant rise in pH from 7.0 to 8.0. Over the next 6 h, there was a steep decline in tetrathionate concentration of the medium, accompanied by an equally steep rise in the concentration of sulfate. After the 18^th^ hour, rate of tetrathionate to sulfate conversion slowed down, and at the 36^th^ hour, 14 mM S tetrathionate was found to remain in the medium together with 26 mM S sulfate (pH of the spent-medium at this time-point was 6.2). Whilst indefinite incubation beyond 36 h resulted in no further oxidation of tetrathionate, re-adjustment of the pH of the spent medium to 8.0 brought about quick oxidation of the residual tetrathionate to sulfate (see graph segments of Fig. 1A that are represented by dotted lines). This result indicated that the tetrathionate-oxidizing machinery of SST functions more efficiently at pH levels above the neutral. The most remarkable aspect of the sulfur oxidation kinetics of *P. thiocyanatus* SST in MST medium was that a small but definite amount of sulfate (total 5 mM S) was produced at a slow rate (0.42 mM S h^-1^) over the first 12 h of incubation when the rest of the 35 mM S thiosulfate (out of the total 40 mM S supplied) was being converted to tetrathionate. In contrast, after thiosulfate completely disappeared form the medium, the rate of sulfate production (now clearly and exclusively from the tetrathionate intermediate formed) became as high as 2.2 mM S h^-1^ between the 12^th^ and 18^th^ hour, and then slowed down to 0.44 mM S h^-1^ between the 18^th^ and 36^th^ hour. These data suggested that the first batch of 5 mM S sulfate potentially came from the direct oxidation of thiosulfate, and the affinity of that thiosulfate oxidation system for its substrate was much lower than the affinity of the thiosulfate-to-tetrathionate converting machinery for the same substrate. It further suggested that the activity of the enzymatic machinery responsible for the oxidation of tetrathionate to sulfate plausibly switches on at ∼pH 8, while its efficiency is reduced at pH 7, and completely limited at pH 6.

**Fig. 1.**
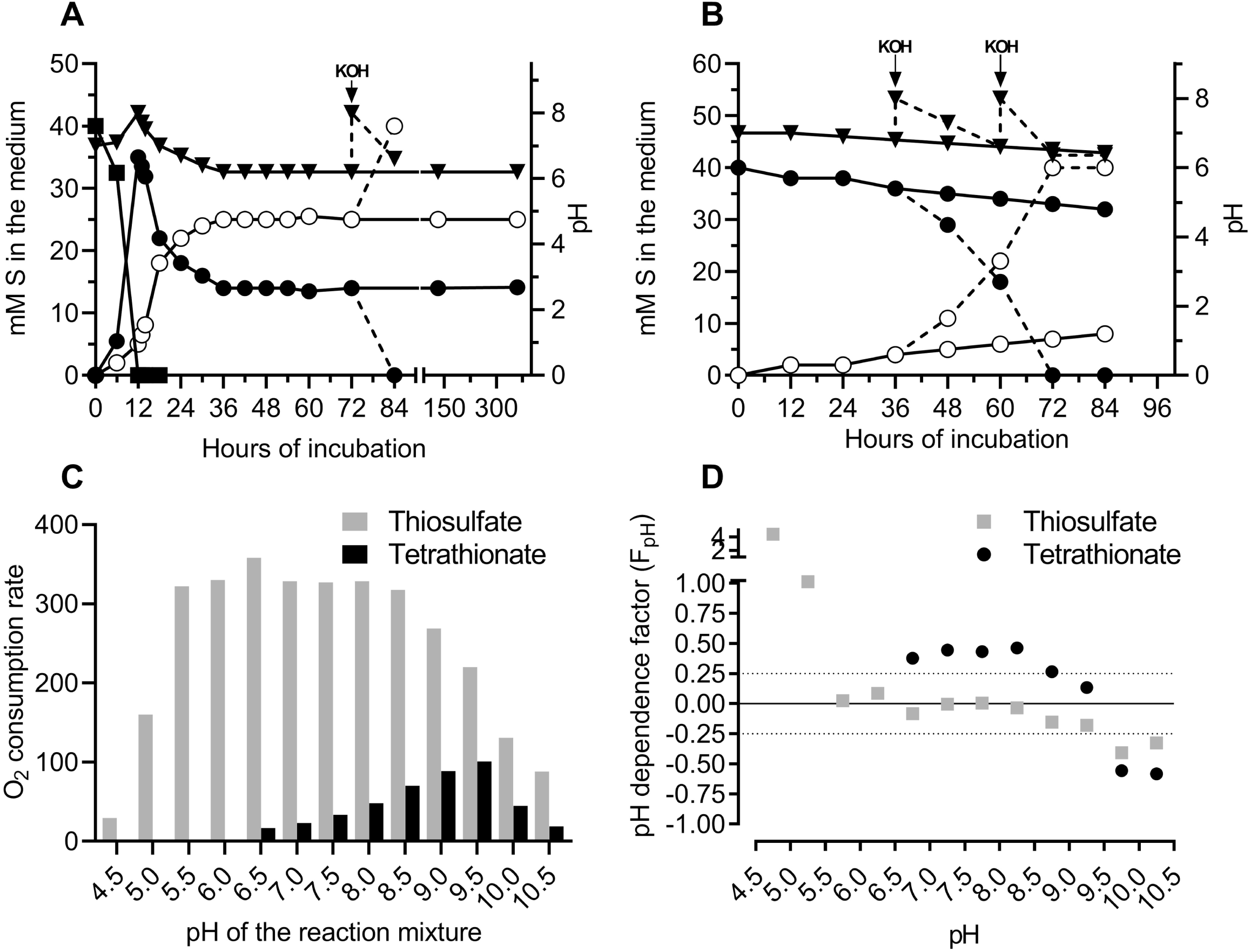
Aspects of thiosulfate and tetrathionate oxidation by *P. thiocyanatus* SST: (**A** and **B**) batch culture kinetics of thiosulfate and tetrathionate oxidation in MST and MSTr media (both having initial pH 7) respectively. Concentrations of S_2_O_3_^2-^ (—▪ —), S_4_O_6_^2-^ (—• —) and SO_4_^2-^ —○ —) in the media are plotted along the primary Y-axes as mM S mL^-1^, while pH values for the media (—▾ —) are in the secondary Y-axes; X-axes represent hours of incubation. Dotted lines indicate separate branches of the experiments where the pH of the media were adjusted twice to 8.0 by addition of extraneous KOH. (**C**) O_2_ consumption rate (in nmol O_2_ consumed mg protein^-1^ min^-1^) of washed SST cells, measured in the presence of thiosulfate (grey bars) or tetrathionate (black bars) at various pH levels. Thiosulfate-and tetrathionate-grown SST cells exhibited almost similar rates of O_2_ consumption in the presence of all sulfur substrates tested, so only the data obtained from thiosulfate-grown cells have been shown. Chemoorganoheterotrophically grown cells, notably, did not exhibit any sulfur-substrate-dependent O_2_ consumption. (**D**) pH dependence factors, calculated (for each pH class at an interval of 0.5) across the pH range 4.5-10.5, for the process of O_2_ consumption by washed SST cells in the presence of thiosulfate (▪) and tetrathionate (•).

### 3.2. pH 9.5 is the optimum for tetrathionate oxidation by SST

When tetrathionate was provided as the initial chemolithotrophic substrate in MSTr medium (initial pH 7.0), *P. thiocyanatus* SST oxidized it very slowly, so after 84 h of incubation in MSTr, only 8 mM S tetrathionate was found to have converted to equivalent amount of sulfate; over this period, pH of the spent medium came down to only 6.4 (Fig. 1B). However, if in the meantime, pH of the spent medium was re-adjusted to 8.0 then the tetrathionate oxidation rate increased remarkably– when this was done at the 36^th^ hour of incubation, the 36 mM S tetrathionate remaining unused at that point of time came down to 18 mM S over the next 24 h (this was accompanied by lowering of pH to 6.6); upon another round of pH-adjustment (to 8.0) at this 60^th^ h, the residual 18 mM S tetrathionate was converted fully to sulfate over the next 12 h (refer to the graph segments of Fig. 1B that are given as dotted lines).

To study the effects of pH on the thiosulfate and tetrathionate oxidation processes of SST, O_2_ consumption rates in the presence of these two sulfur substrates were measured over a pH range of 4.5 to 10.5. Since MST-and MSTr-grown cells exhibited essentially similar rates of O_2_ consumption in the presence of a particular substrate tested (maximum divergence for all data-pairs was ± 5%), data obtained for only the MST grown cells have been presented here. Notably, MSD-grown cells exhibited no O_2_ consumption in the presence of thiosulfate or tetrathionate, thereby indicating that the sulfur oxidation enzyme systems of SST were all induced under chemolithotrophic conditions only.

Thiosulfate-dependent O_2_ consumption rate remained consistently high (i.e. within 317 and 358 nmol O_2_ mg protein^−1^ min^−1^) when pH of the assay buffer was varied between 5.5 and 8.5 (Fig. 1C). Above pH 8.5 O_2_ consumption rate declined and became as low as 88.12 nmol O_2_ mg protein^−1^ min^−1^ at pH 10.5. Highest rate of O_2_ consumption in the presence of tetrathionate (100.82 nmol O_2_ consumed mg protein^−1^ min^−1^) was observed at pH 9.5, which incidentally was higher than the optimum pH (8.0) recorded for SST growth in chemoorganoheterotrophic Luria Bertani medium (Supplementary Table S4). Whereas the tetrathionate-dependent O_2_ consumption rate decreased both below and above the pH 9.5 optima, decline was remarkably sharp between pH 9.5 and 10.5. Values of the pH-dependence factor (*F*_*pH*_) determined for thiosulfate oxidation over the different contiguous 0.5-pH windows remained either zero or very close to zero across the pH range 5.5-8.5. In contrast, very high (4.4) and high (1.0) positive *F*_*pH*_ values were obtained for thiosulfate oxidation over the pH windows 4.5-5.0 and 5.0-5.5 respectively, which indicated that there was a drastic increase in rate of the process as pH changed from 4.5 to 5.0, while the increase in the process rate was also quite sharp with change of pH from 5.0 to 5.5. Gradual decreases in the *F*_*pH*_ values above pH 8.5 indicated a gradual decline in the rate of thiosulfate oxidation with increasing alkalinity (*F*_*pH*_ is slightly < 0.25 for the two windows between pH 8.5 and 9.5, and slightly > 0.25 for the two windows between pH 9.5 and 10.5). While there is no tetrathionate oxidation by strain SST below pH 6.5, *F*_*pH*_ values for were consistently positive, and higher than those obtained for thiosulfate oxidation, over the different pH windows within 6.5-9.5 (Fig. 1D); over the two consecutive windows between pH 9.5 and 10.5, *F*_*pH*_ values were −0.55 and −0.58, numerically higher than the corresponding negative values obtained for thiosulfate oxidation.

### 3.3. Whole genome sequnceingof *P. thiocyanatus* SST and identification of sufur oxidation genes

Shotgun sequencing and annotation of *P. thiocyanatus* SST revealed a 3.53 Mb draft genome having 3,607 potential coding sequences (CDSs). Of all the CDSs identified, 3,285 encoded for proteins. One copy each of the 16S, 23S, and 5S rRNA genes, clustered in a single operon, were also identified together with 45 tRNA genes distributed over the genome. G+C content of the genome was found to be 67.2% (Fig. 2), which is well within the 66-68% G+C content range reported for ∼65% of the genomes sequenced from members of the genus *Paracoccus*. A completeness level of 96.12% was estimated for the assembled draft genome of SST (using the software CheckM) on the basis of the presence of 882 out of a total 917 *Paracoccus*-specific conserved marker genes curated in the CheckM database.

**Fig. 2.**
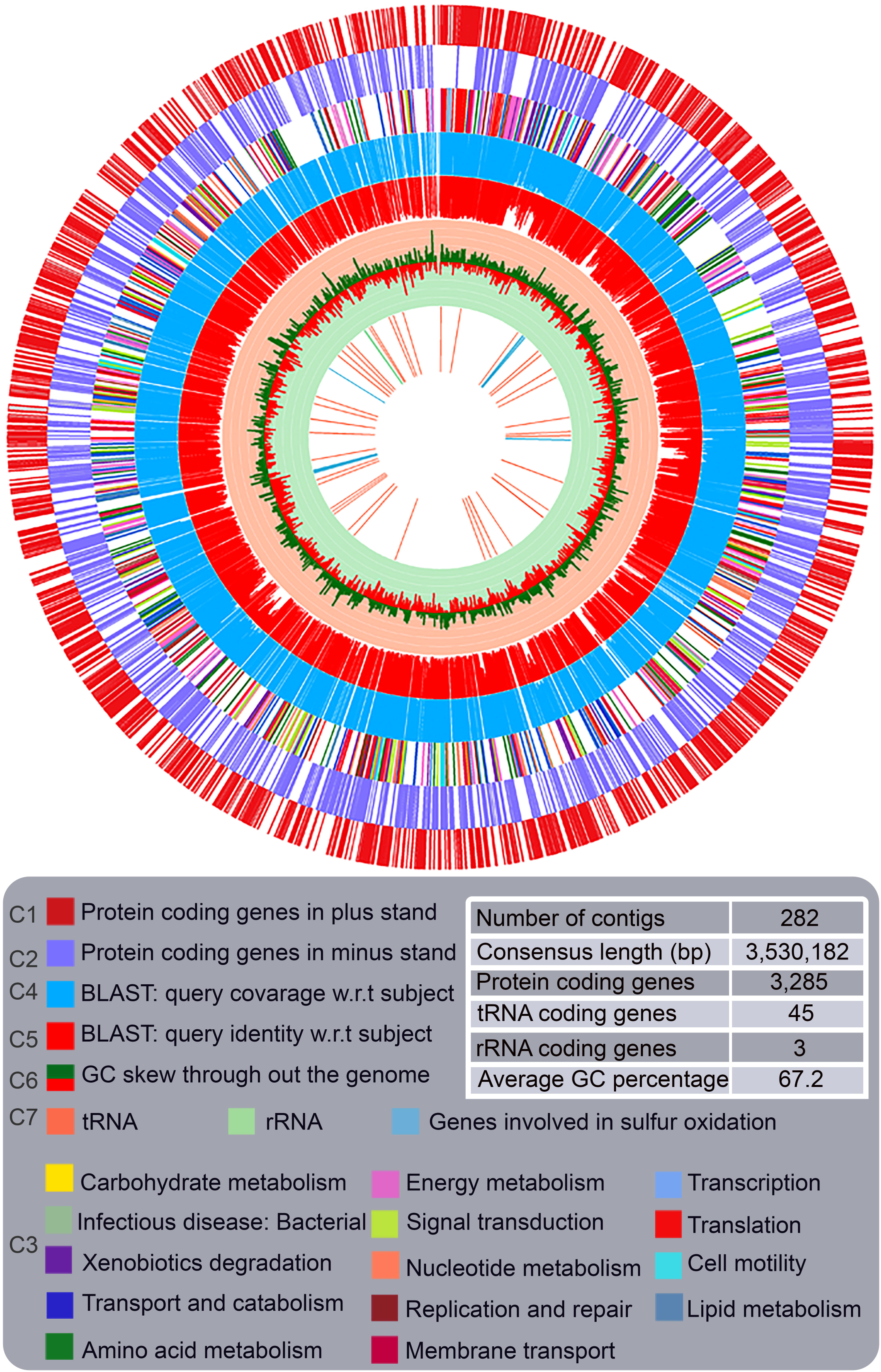
Graphical representation of the assembled and annotated whole genome shotgun sequence of *P. thiocyanatus* SST. The seven concentric circles, from the periphery to center represent the following: (circles 1 and 2) predicted protein-coding regions on the plus and minus strands respectively; (circle 3) protein encoding genes classified into some metabolic categories, based on KEGG orthology (KO) prediction of their products (color-codes for the metabolic categories considered are given in the lower panel); (circles 4 and 5) percentages of coverage and identity obtained respectively for all the protein encoding genes of SST via BlastX search of their translated amino acid sequences against the entire putative protein catalog derived from sequenced *Paracoccus* genomes; (circle 6) GC skew plot embedded in light pink and sea green background; (circle 3) genes encoding homologs of known sulfur oxidation proteins, tRNAs and rRNAs. Order of arrangement, and orientation, of contigs are arbitrary, and so may not tally with the actual genome.

The *P. thiocyanatus* SST genome was found to encompass a complete *soxSRT-VW-XAYZBCD-EFGH* operon similar to the prototypical *sox* operon of *P. pantotrophus* GB17 and *Pseudaminobacter salicylatoxidans* KCT001 in terms of gene synteny (Fig. 3). The putative amino acid sequences of the individual *sox* genes of SST exhibited 74-95% identities with their GB17 counterparts. In addition to the operon-borne *sox* genes, there are duplicate copies of *soxX, soxA, soxC* and *soxD* (locus tags DIE28_02045, DIE28_02050, DIE28_15070 and DIE28_15065 respectively) in the SST genome. These apparently paralogous *sox* gene copies showed 40-62% translated amino acid sequence identities with the operon-coded orthologs of SST/GB17.

**Fig. 3.**
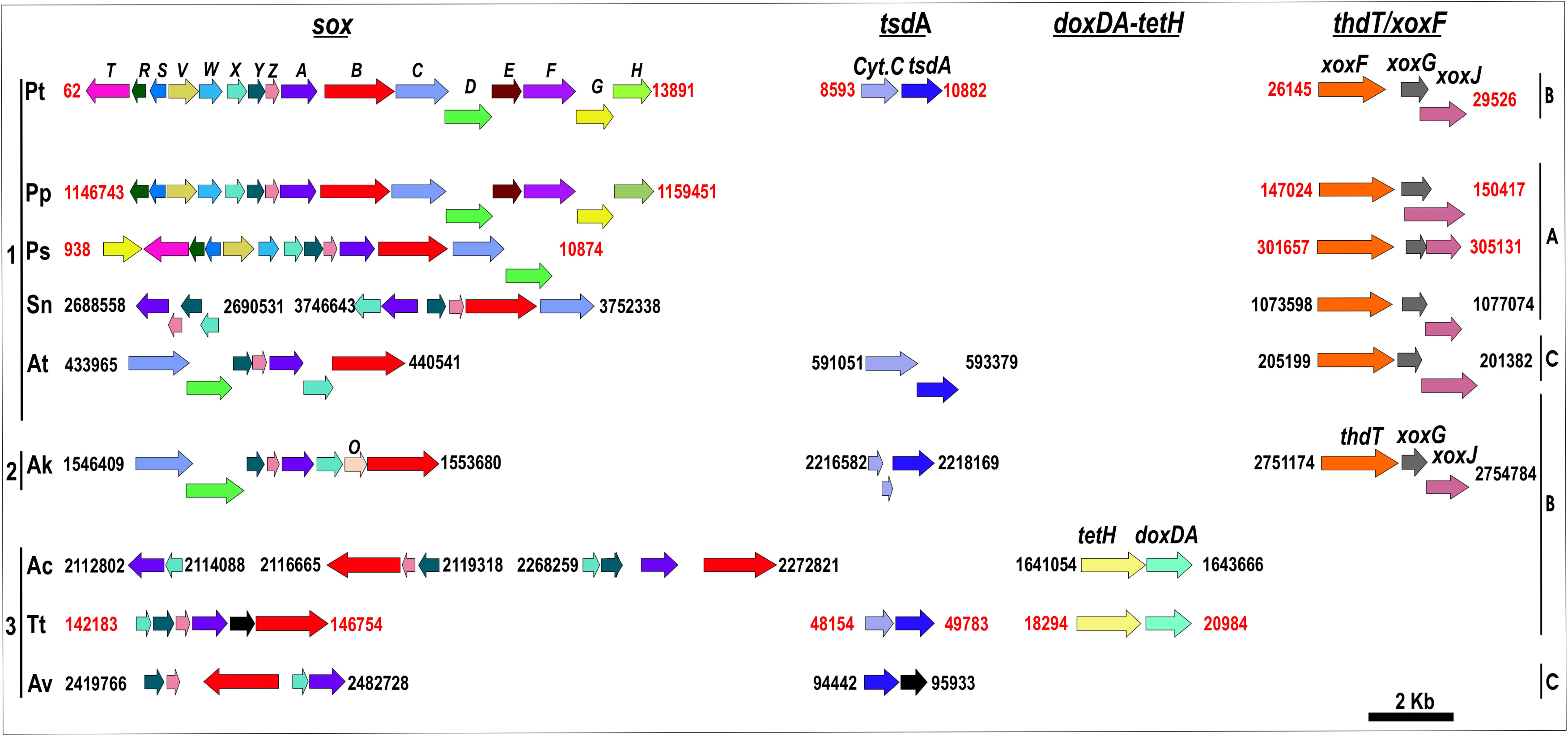
Schematic diagram comparing the sizes and syntenies of the thiosulfate/tetrathionate oxidation genes of *P. thiocyanatus* SST with their homologs or functionally similar counterparts reported from other well-studied sulfur-chemolithotrophs. While the organisms are arranged according to their taxonomy, their respective thiosulfate oxidation phenotypes are also indicated in the right hand side of the figure; so (1, 2 and 3) stand for *Alpha, Beta* and *Gamma* classes of *Proteobacteria*, and (A, B and C) indicate thiosulfate oxidation (using the Sox system) directly to sulfate without any free intermediate, thiosulfate oxidation via S_4_I formation, and conversion of a small portion of thiosulfate to tetrathionate while the rest oxidized to sulfate via other pathways, respectively. Abbreviations used for the different organisms, and the GenBank accession numbers of their genomes or genome contigs where the relevant gene clusters are located, are as follows: **Pt**, *P. thiocyanatus* SST (*sox* - QFCQ01000077, *tsdA*-QFCQ01000015 and *xoxF*-QFCQ01000005); **Pp**, *P. pantotrophus* (*sox*-RBLI01000002.1, *xoxF*-RBLI01000001.1); **Ps**, *P. salicylatoxidans* KCT001 (sox-AJ404005.4, xoxF-NZ_CAIU01000013); **At**, *A. thiophilum* BV-S^T^ (*sox*-NZ_CP012405, *tsdA*-NZ_CP012403, *xoxF*-NZ_CP012405); **Sn**, *Starkeya novella* DSM506^T^ (NC_014217.1); **Tt**, *T.tepidarius* DSM 3134^T^ (*sox*-NZ_AUIS01000007.1, *tsdA*-NZ_AUIS01000009.1, *doxDA*-*tetH*-NZ_AUIS01000032.1); **Ak**, *A. kashmirensis* WT001^T^ (NC_017964.1); **Ac**, *Acidithiobacillus caldus* SM-1 (CP002573.1); and **Av**, *A. vinosum* DSM 180^T^ (CP001896.1). Nucleotide numbers indicate the position of the gene loci on the complete genomes (indicated by black numbers) or partially sequenced genomic segments/contigs (indicated by red numbers) of the organisms that possess them.

A homolog of the *tsd*A gene (locus tag DIE28_04655) was detected in genome of *P. thiocyanatus* SST. The putative amino acid sequence of the protein encoded by this gene was found to encompasse all the functionally-indispensible, and conserved, heme-binding residues characteristic of the prototypical TsdA protein (4WQ7) of *A. vinosum* DSM 180^T^ (Supplementary Fig. S4). In SST, *tsd*A is preceded by a gene (locus tag DIE28_04650) encoding a protein which contains domains similar to those found in the subunit III of di-heme *cbb3* type cytochrome oxidases, designated as *tsdB* in some bacteria (Denkman et al., 2012; Kurth et al., 2016).

Whereas the SST genome annotation did not reveal any tetrathionate hydrolase (*tetH*) gene, three consecutive ORFs (locus tags DIE28_01705, DIE28_01710 and DIE28_01720), homologous to the *xoxF, xoxG* and *xoxJ* genes of the prototypical methanol-oxidizer *Methylobacterium extorquens* AM1 (Nakagawa et al., 2012; Keltjens et al., 2014; Pol et al., 2014; Skovran and Martinez-Gomez, 2015; Chu and Lidstrom, 2016) were detected. Albeit the *xoxF* gene of SST was found to have ∼75% and 69% translated amino acid sequence identities with the typical PQQ-binding XoxF homologs of AM1 and the ThdT variant of XoxF reported from *A. kashmirensis* (Pyne et al., 2018) respectively, the annotated genome sequence of SST deposited in the GenBank designates the locus DIE28_01705 as a pseudogene, apparently due to a frame shift caused by a missing guanine nucleotide in between positions 1161 and 1162 of the ORF. However, a manual scrutiny of the reads aligned to this region of the SST genome showed that 153 and 173 reads taking part in the assembly contained and lacked the guanine in the above mentioned position respectively, while 36 others had an ambiguous base call in this nucleotide position (notably, proportion of ambiguous base calls across this *xoxF* ORF was also quite high). As capillary sequencing also failed to resolve this issue unambiguously, we decided to knock this gene out and confirm whether tetrathionate oxidation in SST proceeded via a ThdT-mediated glutathione-coupled mechanism similar to the one described for *A. kashmirensis* (Pyne et al., 2017, 2018). Other sulfur-oxidation-related genes (see Ghosh and Dam, 2009, and references therein), such as those encoding flavocytochrome *c*-sulfide dehydrogenase (FCSD), sulfide:quinine oxidoreductase (SQOR), sulfite:acceptor oxidoreductase (SorAB), and the reverse-acting dissimilatory sulfite reductase (rDsr complex), were not detected in the SST genome.

So far as the genomic basis of thiocyanate oxidation by SST is concerned, an ORF (locus tag DIE28_02005) whose putative protein product exhibits 36-86% identities withthe well-studied thiocyanate dehydrogenase (TcDH, Protein Database Accession: 5F75) of *Thioalkalivibrio, Thiohalobacter* and *Guyparkeria* species (Berben et al., 2017; Tsallagov et al., 2019) was detected in the genome of this aphaproteobacterium. TcDH has been hypothesized to attack the sulfane sulfur atom of thiocyanate (SCN^-^) and directly oxidize SCN^-^ to elemental sulfur (S^0^) and cyanate (CNO^-^) (Sorokin et al., 2001; Tsallagov et al., 2019). If thiocyanate oxidation in SST proceeds via this pathway, then the sulfur atom produced can be oxidized via the Sox machinery encoded by the genome of this alphaproteobacterium; notably however, no homolog of the cyanate hydratase (cyanase) gene, which is known to convert cyanate to ammonia (NH_3_) and carbon dioxide (CO_2_) (Anderson and Little, 1986; Sung and Fuchs, 1988), was detected in the SST genome. It, therefore, seems plausible that there is some hitherto unknown mechanism in SST to take care of the CNO^-^ radical. In relation to thiocynate oxidation it is further noteworthy that annotation of the SST genome did not reveal any genes for the three subunits of thiocyanate hydrolase, which degrades thiocyanate via the carbonyl sulfide (COS) pathway (i.e. conversion of thiocyanate to COS and NH_3_) in organisms such as *Thiobacillus thioparus* and *Thiohalophilus thiocyanoxidans* (Katayama et al., 1998; Bezsudnova et al., 2007).

### 3.4. Knock-outmutantion of tsdA abolishes S_4_I formation but thiosulfate is still oxidized to sulfate without any free intermediate

The Δ*tsdA* knock-out mutant of *P. thiocyanatus* SST (Fig. 4A), designated as Pt_KoT, when grown in MST medium (initial pH 7.0), did not convert any thiosulfate to tetrathionate (Fig. 4B), thereby confirming that the *tsdA* is the only gene responsible for this metabolic transformation in SST. Remarkably, however, Pt_KoT converted 26 mM S thiosulfate to sulfate without the formation of any free intermediate, over 210 h incubation in MST (Fig. 4B). Tetrathionate oxidation properties of Pt_KoT in MSTr medium (initial pH 8.0) were similar to those of the wild-type SST strain (Fig. 4C). Whilst this thiosulfate to sulfate oxidation phenotype of Pt_KoT was reflective of the potential involvement of the Sox pathway, the total amount of sulfate (5 mM S) produced by this mutant strain over the first 12 h of its incubation in MST was exactly same as that produced by the wild-type over the same period of incubation in the same medium. Furthermore, the rate of the process in Pt_KoT during the exponential phase of sulfate production, i.e., between the 6^th^ and the 30^th^ hours of growth in MST (this was calculated to be 0.6 mM S h^-1^; see Fig. 4B), was essentially similar to the rate of sulfate production observed for the wild-type strain between the 6^th^ and the 12^th^ hours of its growth in MST, i.e. during the brief exponential phase of the first stage of sulfate productionby SST (this was calculated to be 0.5 mM S h^-1^; see Fig. 1A). These data indicated that the first batch of small amount of sulfate (5 mM S) produced by SST over the first 12 h of its incubation in MST medium came from came directly from thiosulfate and the S_4_I had no contribution in its formation.

**Fig. 4.**
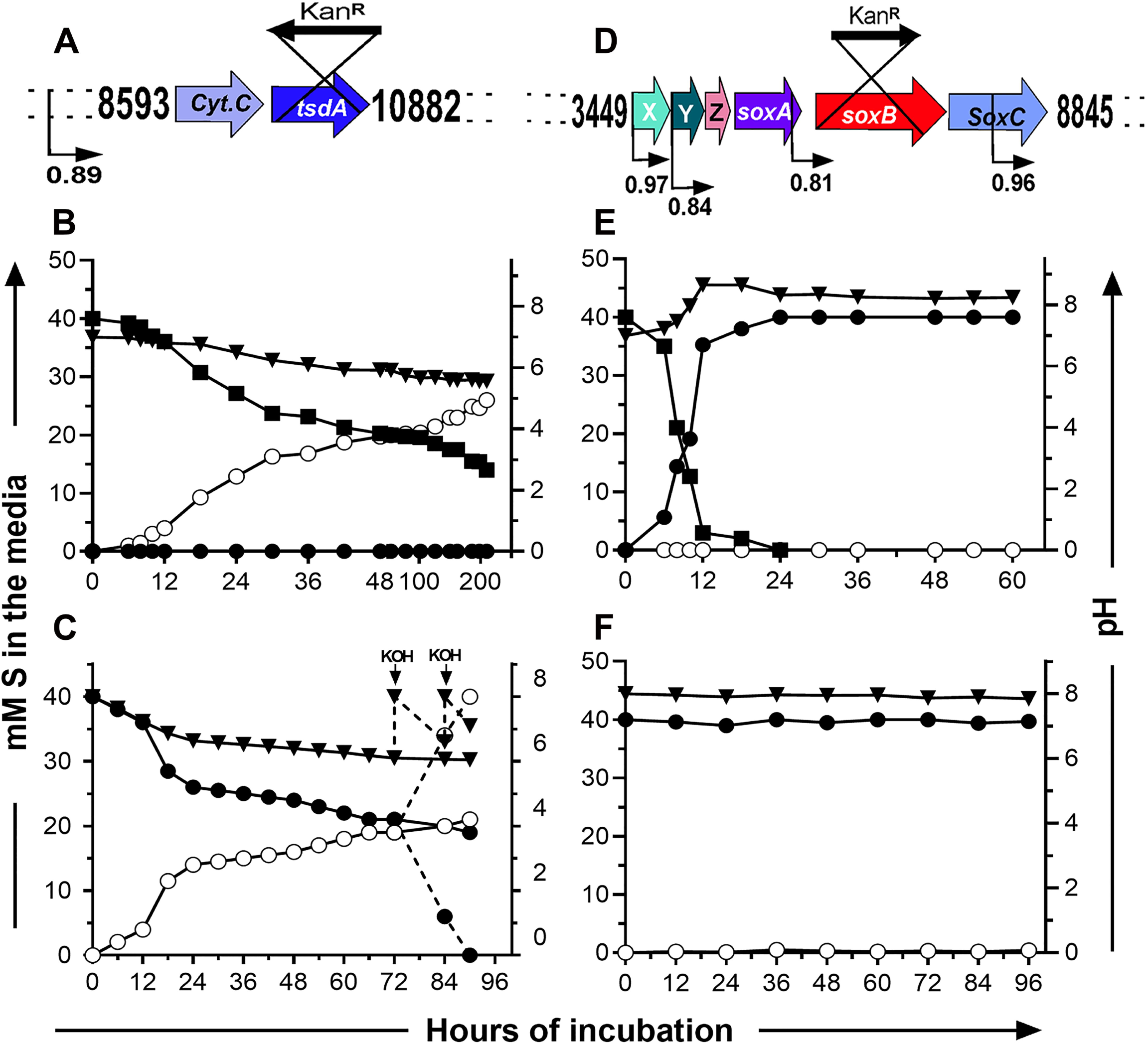
Homologous-recombination-based knock-out mutation of the *tsdA* and *soxB* genes of SST, and test of the sulfur-oxidation phenotypes of the resultant mutants. (**A**) genomic context of the *tsdA* mutation; (**B** and **C**) batch culture kinetics of thiosulfate and tetrathionate oxidation by Pt_KoT in MST and MSTr media (initial pH 7.0 and 8.0) respectively. Dotted lines in (**C**) indicate a separate branch of the experiment where the pH of the medium was adjusted twice to 8.0 by addition of extraneous KOH. (**D**) genomic context of the *soxB* mutation; (**E** and **F**) batch culture kinetics of thiosulfate and tetrathionate oxidation by Pt_KoB in MST and MSTr media (initial pH 7.0 and 8.0) respectively. In (**A** and **D**) promotors and their corresponding prediction probability values are indicated by bent arrows and numbers below them respectively. In (**B** and **C**, and **E** and **F**) concentrations of S_2_O_3_^2-^ (—▪ —), S_4_O_6_^2-^ (—•—) and SO_4_^2-^ (—o—) in the media are plotted along the primary Y-axes as mM S mL^-1^, while pH values for the media (—▾—) are in the secondary Y-axes; X-axes represent hours of incubation.

### 3.5. Knock-out mutantion of SST *soxB* abolished the production of sulfate from thiosulfate as well as tetrathionate

The Δ*soxB* knock-out mutant of *P. thiocyanatus* SST (Fig. 4D), designated as strain Pt_KoB, when grown in MST (initial pH 8.0) medium, did not produce even the slightest of sulfate (Fig. 4E), whereas the entire 40 mM S thiosulfate was converted to tetrathionate over 24 h incubation, with concomitant increase in the pH of the spent medium to 8.7 (Fig. 4E). The S_4_I formed, however, remained unutilized even after prolonged incubation of Pt_KoB in this medium. These data confirmed that the 5 mM S sulfate that appeared in the first 12 h incubation of wild-type SST in the MST medium indeed came from thiosulfate via the Sox pathway. Pt_KoB was also unable to oxidize any tetrathionate when grown in MSTr medium (initial pH 8.0) corroborated by no sulfate production (Fig. 4F), thereby indicating the indispensibility of SoxB in tetrathionate oxidation by *P. thiocyanatus*.

### 3.6. Tetrathionate oxidation by *P. thiocyanatus* involves thiol groups, particularly glutathione

Involvement of thiol groups in the sulfur oxidation mechanism of *P. thiocyanatus* was tested by measuring the thiosulfate and tetrathionate-dependent O_2_ consumption rates of the washed cells of the wild-type SST, as well as those of its mutants, pre-treated/untreated with the thiol-binding reagent N-ethyl maleimide (NEM) and the thiol group containing tripeptide glutathione (GSH).

For the wild-type SST, thiosulfate-dependent O_2_ consumption rate was reduced by approximately 17% when cells were pre-treated with NEM; pre-treatment with GSH however had no effect on the thiosulfate-dependent O_2_ consumption rate of SST (Fig. 5A). In contrast, O_2_ consumption rate of the NEM-treated SST cells in the presence of tetrathionate (15.91nmol O_2_ mg protein^-1^ min^-1^) was ∼70% less than that of their untreated counterparts [51.07 nmol O_2_ mg protein^-1^ min^-1^(Fig. 5B)]. When the washed cells of SST were pre-treated with 10 mM GSH, tetrathionate-dependent O_2_ consumption rate (136.22 nmol O_2_ mg protein^-1^ min^-1^) increased by 2.7 folds, as compared to that of the untreated cells (Fig. 5B). The wild-type SST showed no O_2_ consumption in the presence of sulfite, so was not tested for this parameter in the presence of NEM/GSH (Fig. 5C). Notably, no consumption of O_2_ in the presence of sulfite by the wild-type SST cells was consistent with consistent with the fact that no free sulfite was detected during tetrathionate oxidation by SST and also that no *sorAB* gene was detected in annotated genome of this organism.

**Fig. 5.**
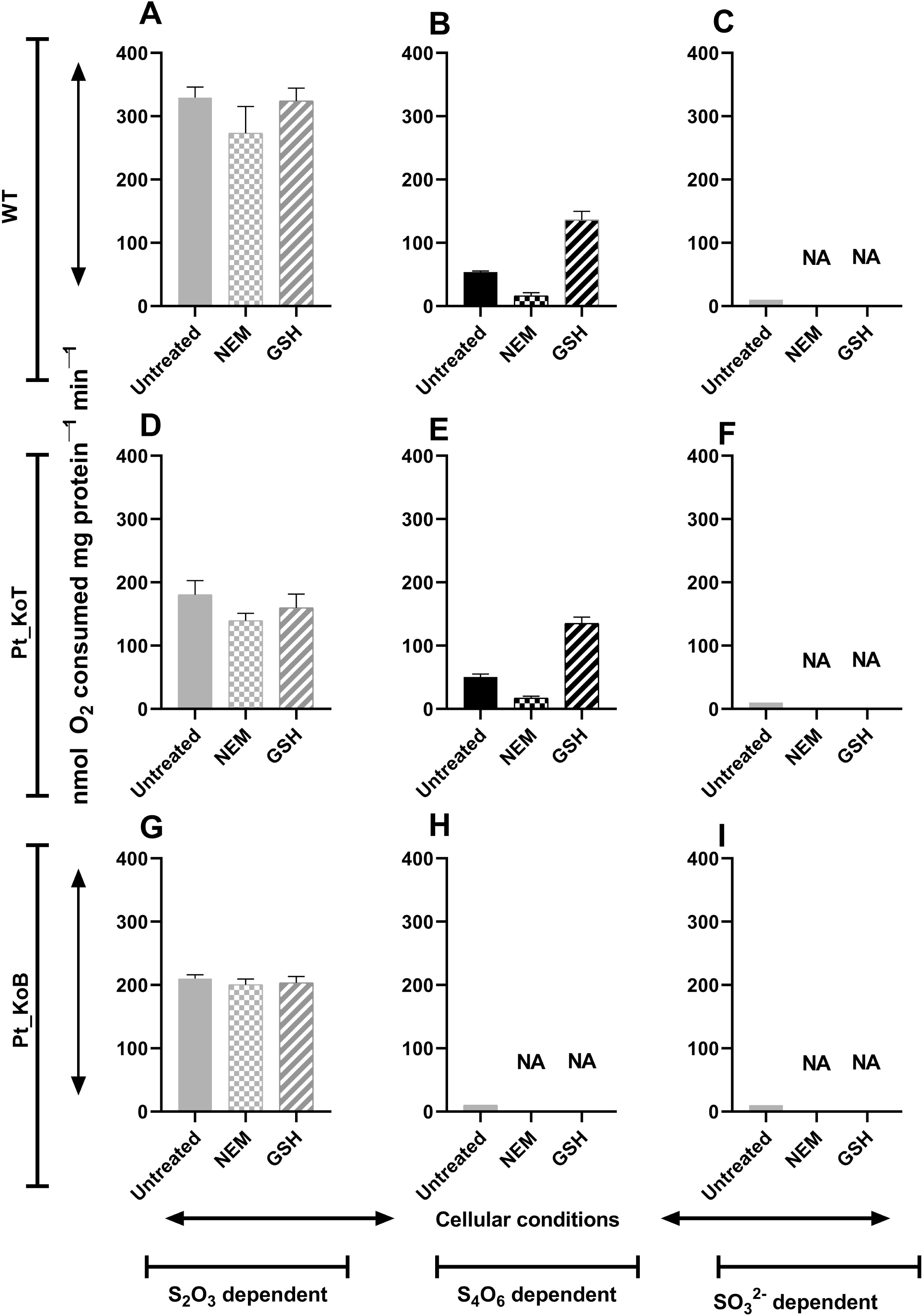
Effect of GSH and NEM on sulfur-substrate-dependent O_2_ consumption rates (in nmol O_2_ consumed mg protein^-1^ min^-1^) of washed whole cells of *P. thiocyanatus* SST and its mutants, untreated or pre-treated with NEM and GSH. (**A, B**, and **C**) O_2_ consumption in the presence of thiosulfate, tetrathionate and sulfite by the wild type respectively; (**D, E** and **F**) O_2_ consumption in the presence of thiosulfate, tetrathionate and sulfite by Pt_KoT respectively; (**G, H** and **I**) O_2_ consumption in the presence of thiosulfate, tetrathionate and sulfite by Pt_KoB respectively. Since thiosulfate- and tetrathionate-grown cells of all three strains showed similar rate of O_2_ consumption in the presence of thiosulfate, tetrathionate and sulfite, data for only the MST-grown cells have been shown.

For the Δ*tsdA* mutant Pt_KoT, thiosulfate-dependent O_2_ consumption rates of the untreated, NEM-treated and GSH-treated cells (Fig. 5D) were all more or less reduced by 40-50% from the corresponding wild-type levels (Fig. 5A). Since thiosulfate oxidation in the mutant Pt_KoT did not proceed via S_4_I at all, and instead produced sulfate directly from the substrate, these data clearly corroborated the slower pace of the Sox-based thiosulfate oxidation process of *P. thiocyanatus*, as compared to its S_4_I Pathway through which almost 90% of the thiosulfate passes during oxidation by the wild-type. Expectedly, tetrathionate- or sulfite-dependent O_2_ consumption phenotypes of untreated / NEM-treated / GSH-treated Pt_KoT cells (Fig. 5E-F), were same as the corresponding wild-type phenotypes (Fig. 5B-C).

For the Δ*soxB* mutant Pt_KoB, which only converted thiosulfate to tetrathionate and could not oxidize the latter, thiosulfate-dependent O_2_ consumption rate of untreated cells was ∼64% of the corresponding wild-type level; NEM-/GSH-treatment did not affect this thiosulfate-dependent O_2_ consumption rate (Fig. 5G). Even the NEM-/GSH-untreated cells of this mutant had no tetrathionate- or sulfite-dependent O_2_ consumption phenotype (Fig. 5H-I), reiterating the involvement of SoxB in SST tetrathionate oxidation.

In view of the above indications for the involvement of glutathione in the tetrathionate oxidation mechanism of *P. thiocyanatus* SST, it was imperative to check whether this involved any XoxF-homolog-mediated tetrathionate-glutathione coupling, which apparently results in the production of a glutathione:sulfodisulfane adduct and sulfite in the betaproteobacterium *A. kashmirensis* (Pyne et al., 2017, 2018). The *xoxF* homolog (locus tag DIE28_01705) present in the *P. thiocyanatus* SST genome was, therefore, inactivated by knock-out mutation (Fig. 6A). The resultant mutant Pt_KoX, interestingly, exhibited wild-type-like phenotype during chemolithoautotrophic growth in both MST as well as MSTr media (Fig. 6B-C). These results implied that the involvement of glutathione in the SST tetrathionate oxidation mechanism is not ThdT-/XoxF-mediated, and therefore may or may not initialize via formation of a glutathione:sulfodisulfane adduct and sulfite.

**Fig. 6.**
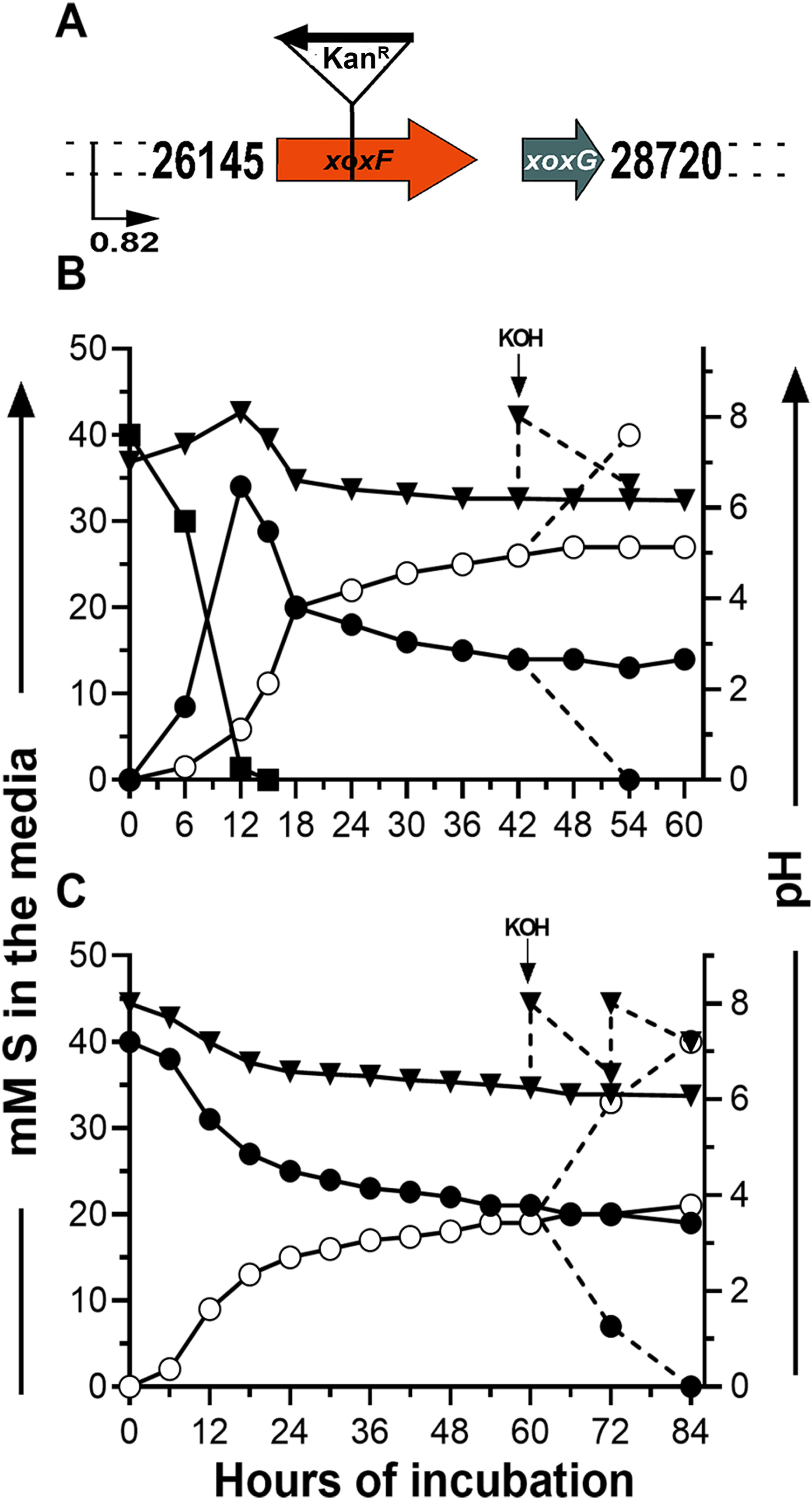
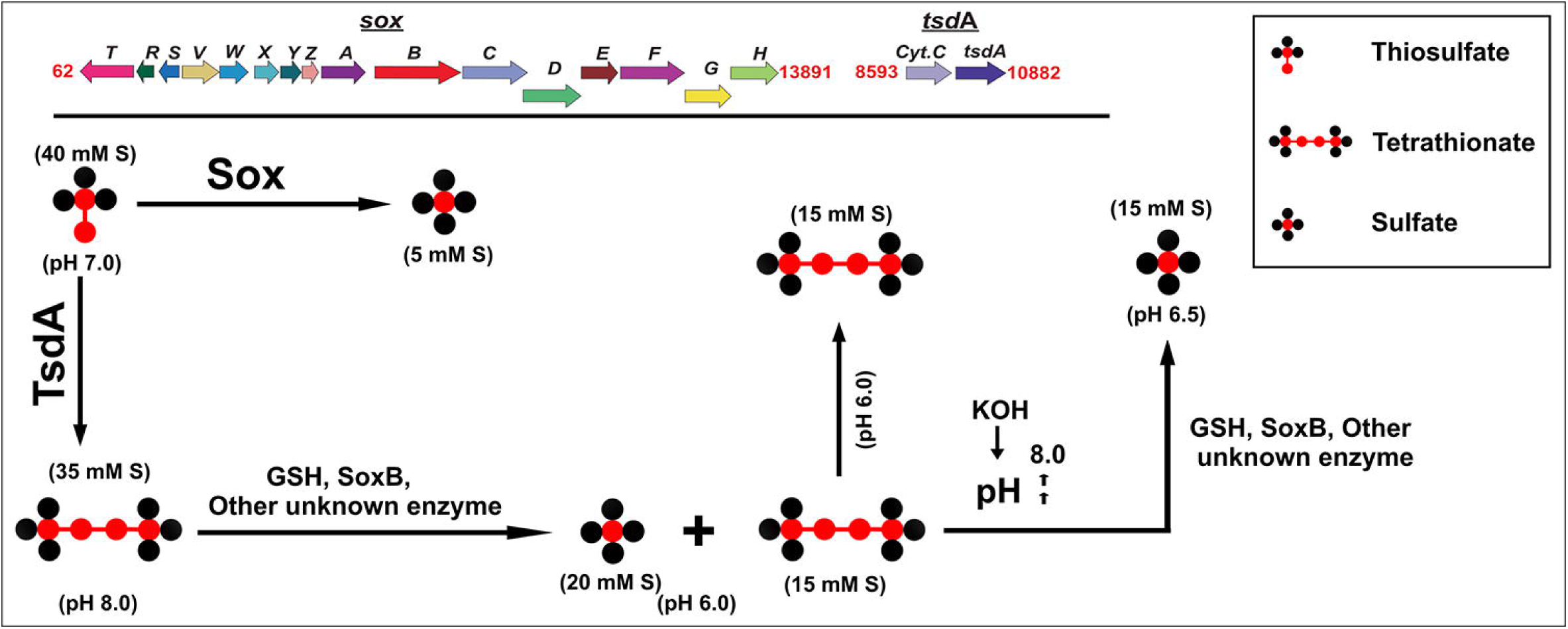
Homologous-recombination-based knock-out mutation of the *xoxF* gene of SST, and test of the sulfur-oxidation phenotypes of the resultant mutant. (**A**) genomic context of the *xoxF* mutation; position of the promotor and its corresponding prediction probability value are indicated by a bent arrow and the numbers below it respectively. (**B** and **C**) batch culture kinetics of thiosulfate and tetrathionate oxidation by Pt_KoX in MST and MSTr media (both having initial pH 7) respectively. Dotted lines indicate separate branches of the experiments where the pH of the media were adjusted twice to 8.0 by addition of extraneous KOH. Concentrations of S_2_O_3_^2-^ (—▪—), S_4_O_6_^2-^ (—•—) and SO_4_^2-^ (—o—) in the media are plotted along the primary Y-axes as mM S mL^-1^, while pH values for the media (—▾ —) are in the secondary Y-axes; X-axes represent hours of incubation.

### 3.7. Novel aspects of the thiosulfate and tetrathionate oxidation pathways of *P. thiocyanatus* SST

Summing up of all the physiological, genomic, and molecular genetic data obtained under the current investigation, revealed the following.

- Formation and subsequent oxidation of the S_4_I is the main pathway of thiosulfate oxidation in *P. thiocyanatus* SST.
- During thiosulfate oxidation, tetrathionate as well as sulfate formation starts together under the mediation of the TsdA and Sox systems respectively; but the former process far out-competes the latter in terms of the kinetics of sulfur atoms oxidized over time.
- Oxidation of tetrathionate, whether produced from thiosulfate or supplied as a starting substrate for chemolithotrophic growth, is a highly pH-sensitive process having its optima at pH 9.5.
- Tetrathionate oxidation in SST involves a clear role of glutathione, even though the mechanistic basis of this involvement is apparently distinct from the ThdT-mediated glutathione:sulfodisulfane and sulfite formation typical of *A. kashmirensis*.
- Some hitherto unknown enzyme(s) seems to catalyze glutathione:tetrathionate interaction in SST. In such a scenario the typical thiol esterase and sulfur dehydrogenase roles of SoxB and SoxCD respectively (Friedrich et al., 2001, 2005), which have been envisaged for the oxidation of glutathione:sulfodisulfane in *A. kashmirensis* (Pyne et. al., 2018), may also not be there in SST.

The substantial body of knowledge currently generated on the thiosulfate- and tetrathionate-based chemolithotrophy of *P. thiocyanatus* SST points out a number of novel facts regarding the mechanistic diversification of sulfur-chemolithrophy. Formation, accumulation and subsequent oxidation of S_4_I as the primary mechanism of chemolithotrophic thiosulfate oxidation in a member of *Paracoccus* (despite a fully functional Sox system being in place) is surprising as it is idiosyncratic to the established preeminence of the typical Sox pathway in thiosulfate oxidation across *Alphaproteobacteria*, and especially the genus *Paracoccus* from where most of the information on Sox emanated. So we believe that the findings of this study would shed new light on our understanding of mechanistic diversification of sulfur-chemolithotrophy not only in phylogenetically unrelated but also in closely-related bacteria as well.

The domination of the S_4_I pathway over Sox in *P. thiocyanatus* SST is clearly attributable to the presence of a very strong TsdA which not only has the ability to convert high quantities of thiosulfate to tetrathionate, but also is empowered with very high affinity for thiosulfate. SST TsdA’s affinity towards thiosulfate could be 7 times more than that of the SST Sox system – this is apparent from the fact that while growing the wild-type strain in MST, out of 40 mM S thiosulfate provided, 35 mM S was converted to tetrathionate and only 5 mM S got converted to sulfate, apparently via Sox, over the first 12 h of incubation. At the same time, it is peculiar that the Sox system of SST is far more sluggish than that of *Paracoccus pantotrophus*; for instance, in the same MST medium, we had previously reported *P. pantotrophus* strain LMG 4218 to oxidize approximately 85% of the supplied 40 mM S thiosulfate directly to sulfate over an incubation period of 48 h (Alam et al., 2013). Furthermore, the heavy-duty feat of SST TsdA is clearly evident when its present activity is compared with that of the other TsdA homologs, well-studied for their activites. For instance, the SoxCD-less photolithoautotroph *A.vinosum* DSM 180^T^, at pH < 7.0, forms only 0.4 mM S tetrathionate from the total 5 mM S thiosulfate supplied, by the action of its very well-studied TsdA (Hensen et al., 2006; Brito et al., 2014; 2015; Kurth et al., 2016) whereas the rest of the thiosulfate is oxidized to sulfate, via formation of elemental sulfur, by the step-wise action of SoxXAYZB and the rDsr system (Hensen et. al., 2006). Similarly, in *Thiomicrospira thermophila* EPR85, thiosulfate is oxidized to sulfate with formation of extracellular elemental sulfur at pH 5.5-8.0; but at pH < 5.5, a fraction of thiosulfate is converted to tetrathionate with a sulfate:tetrathionate formation ratio of 1:0.46 (Houghton et. al., 2016).

Whilst a new generation of novel gene search via transcriptomic, proteomic and/or metabolomic studies would, in the near future, track down the unknown enzyme(s) governing glutathione:tetrathionate interaction in SST, the findings of the present paper illuminate broader aspects of sulfur chemolihtotrophy, such as the mechanistic diversification of sulfur-chemolithotrophy in phylogenetically related and unrelated bacteria. Furthermore, continued biochemical and biophysical investigations with TsdA and Sox in tandem can open up new vistas of understanding on how an enzyme’s/enzyme system’s biochemical vigor for its typical activity varies across the plethora of homologs. This, in turn, can become one of the key determinants of whether, in practice, a homolog at all manages to obtain any employment within the highly-competitive job market of a microbial cell.

## Acknowledgements

This research was funded by Bose Institute, Department of Science and Technology, Government of India (GOI). MJR recieved doctoral fellowship from University Grants Commission, GOI. PP and JS received fellowship from the Council of Scientific and Industrial Research (CSIR), GOI. SM received doctoral fellowship from Department of Science and Technology, GOI. SB received doctoral fellowship from Bose Institute. NM received doctoral fellowship from Science and engineering research board, GOI.

## Appendix A. Supplementary data

Supplemental data for this article may be found with the digital version of this manuscript.

## References

Alam, M., Pyne, P., Mazumdar, A., Peketi, A., Ghosh, W., 2013. Kinetic enrichment of ^34^S during proteobacterial thiosulfate oxidation and the conserved role of SoxB in S-S bond breaking. Appl. Environ. Microbiol., 79, 4455–4464. https://doi.org/10.1128/AEM.00956-13.

Anderson, P.M., Little, R.M., 1986. Kinetic properties of cyanase. Biochemistry 25, 1621–1626. https://doi.org/10.1021/bi00355a026.

Appia-Ayme, C., Little, P.J., Matsumoto, Y., Leech, A.P., Berks, B.C., 2001. Cytochrome complex essential for photosynthetic oxidation of both thiosulfate and sulfide in *Rhodovulum sulfidophilum*. J. Bacteriol. 183, 6107–6118. https://doi.org/10.1128/JB.183.20.6107-6118.2001.

Bamford, V.A., Bruno, S., Rasmussen, T., Appia-Ayme, C., Cheesman, M.R., Berks, B.C., Hemmings, A.M., 2002. Structural basis for the oxidation of thiosulfate by a sulfur cycle enzyme. EMBO J. 21, 5599–5610. https://doi.org/10.1093/emboj/cdf566.

Bankevich, A., Nurk, S., Antipov, D., Gurevich, A.A., Dvorkin, M., Kulikov, A.S., Lesin, V.M., Nikolenko, S.I., Pham, S., Prjibelski, A.D., Pyshkin, A.V., Sirotkin, A.V., Vyahhi, N., Tesler, G., Alekseyev, M.A., Pevzner, P.A., 2012. SPAdes: a new genome assembly algorithm and its applications to single-cell sequencing. J. Comput. Biol. 19, 455–477. https://doi.org/10.1089/cmb.2012.0021.

Berben, T., Overmars, L., Sorokin, D.Y., Muyzer, G., 2017. Comparative genome analysis of three thiocyanate oxidizing *Thioalkalivibrio* species isolated from soda lakes. Front Microbiol, 8, Article 254. https://doi.org/10.3389/fmicb.2017.00254.

Bezsudnova, E.Y., Sorokin, D.Y., Tikhonova, T.V., Popov, V.O., 2007. Thiocyanate hydrolase, the primary enzyme initiating thiocyanate degradation in the novel obligately chemolithoautotrophic halophilic sulfur-oxidizing bacterium *Thiohalophilus thiocyanoxidans*. Biochim. Biophys. Acta 1774, 1563–1570. https://doi.org/10.1016/j.bbapap.2007.09.003.

Brito, J.A., Gutierres, A., Denkmann, K., Dahl, C., Archer, M., 2014. Production, crystallization and preliminary crystallographic analysis of *Allochromatium vinosum* thiosulfate dehydrogenase TsdA, an unusual acidophilic c-type cytochrome. Acta Cryst. F70, 1424–1427. https://doi.org/10.1107/S2053230X14019384.

Brito, J.A., Denkmann, K., Pereira, Inês.A.C., Archer, M., Dahl, C., 2015. Thiosulfate Dehydrogenase (TsdA) from *Allochromatium vinosum*:structural and functional insights into thiosulfate oxidation. J. Biol. Chem. 290, 9222–9238. https://doi.org/10.1074/jbc.M114.623397.

Bugaytsova, Z., Lindström, E.B., 2004. Localization, purification and properties of a tetrathionate hydrolase from *Acidithiobacillus caldus*. Eur. J. Biochem. 271, 272–280.https://doi.org/10.1046/j.1432-1033.2003.03926.x.

Chu, F., Lidstrom, M.E., 2016. XoxF acts as the predominant methanol dehydrogenase in the type I methanotroph *Methylomicrobium buryatense*. J. Bacteriol. 198, 1317–1325. https://doi.org/10.1128/JB.00959-15.

Cutter, G.A., Kluckhohn, R.S., 1999. The cycling of particulate carbon, nitrogen, sulfur, and sulfur species (iron monosulfide, greigite, pyrite, and organic sulfur) in the water columns of Framvaren Fjord and the Black Sea. Mar. Chem. 67, 149–160. https://doi.org/10.1016/S0304-4203(99)00056-0.

Dahl, C., 2008. Inorganic Sulfur Compounds as Electron Donors in Purple Sulfur Bacteria, in: Hell, R., Dahl, C., Knaff, D., Leustek, T. (Eds.), Sulfur Metabolism in Phototrophic Organisms. Advances in Photosynthesis and Respiration,. Springer, New York, pp. 289–317. https://doi.org/10.1007/978-1-4020-6863-8_15.

Dam, B., Mandal, S., Ghosh, W., Gupta, S.K.D., Roy, P., 2007. The S4-intermediate pathway for the oxidation of thiosulfate by the chemolithoautotroph *Tetrathiobacter kashmirensis* and inhibition of tetrathionate oxidation by sulfite. Res. Microbiol. 158, 330–338. https://doi.org/10.1016/j.resmic.2006.12.013.

De Jong, G.A.H., Hazeu, W., Bos, P., Kuenen, G., 1997a. Polythionate degradation by tetrathionate hydrolase of *Thiobacillus ferrooxidans*. Microbiology 143, 499–504. https://doi.org/10.1099/00221287-143-2-499.

De Jong, G.A.H., Hazeu, W., Bos, P., Kuenen, J.G., 1997b. Isolation of the tetrathionate hydrolase from *Thiobacillus acidophilus*. Eur. J. Biochem. 243, 678–683. https://doi.org/10.1111/j.1432-1033.1997.00678.x.

Denkmann, K., Grein, F., Zigann, R., Siemen, A., Bergmann, J., van Helmont, S., Nicolai, A., Pereira, I.A.C., Dahl, C., 2012. Thiosulfate dehydrogenase: a widespread unusual acidophilic c-type cytochrome. Environ. Microbiol. 14, 2673–2688. https://doi.org/10.1111/j.1462-2920.2012.02820.x.

Eisen, J.A., Nelson, K.E., Paulsen, I.T., Heidelberg, J.F., Wu, M., Dodson, R.J., Deboy, R., Gwinn, M.L., Nelson, W.C., Haft, D.H., Hickey, E.K., Peterson, J.D., Durkin, A.S., Kolonay, J.L., Yang, F., Holt, I., Umayam, L.A., Mason, T., Brenner, M., Shea, T.P., Parksey, D., Nierman, W.C., Feldblyum, T.V., Hansen, C.L., Craven, M.B., Radune, D., Vamathevan, J., Khouri, H., White, O., Gruber, T.M., Ketchum, K.A., Venter, J.C., Tettelin, H., Bryant, D.A., Fraser, C.M., 2002. The complete genome sequence of *Chlorobium tepidum* TLS, a photosynthetic, anaerobic, green-sulfur bacterium. Proc. Natl. Acad. Sci. USA. 99, 9509–14. https://doi.org/10.1073/pnas.132181499.

Friedrich, C.G., Quentmeier, A., Bardischewsky, F., Rother, D., Kraft, R., Kostka, S., Prinz, H., 2000. Novel Genes Coding for Lithotrophic Sulfur Oxidation of *Paracoccus pantotrophus* GB17. J. Bacteriol. 182, 4677–4687. https://doi.org/10.1128/JB.182.17.4677-4687.2000.

Friedrich, C.G., Rother, D., Bardischewsky, F., Quentmeier, A., Fischer, J., 2001. Oxidation of reduced inorganic sulfur compounds by bacteria, emergence of a common mechanism? Appl. Environ. Microbiol. 67, 2873–2882. https://doi.org/10.1128/AEM.67.7.2873-2882.2001.

Friedrich, G., Bardischewsky, F., Rother, D., Quentmeier, A., Fischer, J., 2005. Prokaryotic sulfur oxidation. Curr. Opin. Microbiol. 8, 253–259. https://doi.org/10.1016/j.mib.2005.04.005.

Frolov, E.N., Belousova, E.V., Lavrinenko, K.S., Dubinina, G.A., Grabovich, M.Yu., 2013. Capacity of *Azospirillum thiophilum* for lithotrophic growth coupled to oxidation of reduced sulfur compounds. Microbiology 82, 271–279. https://doi.org/10.1134/S0026261713030053.

Ghosh, W., Bagchi, A., Mandal, S., Dam, B., Roy, P., 2005. *Tetrathiobacter kashmirensis* gen. nov., sp. nov., a novel mesophilic, neutrophilic, tetrathionate-oxidizing, facultatively chemolithotrophic betaproteobacterium isolated from soil from a temperate orchard in Jammu and Kashmir, India. Int. J. Syst. Evol. Microbiol. 55, 1779–1787. https://doi.org/10.1099/ijs.0.63595-0.

Ghosh, W., Roy, P., 2007. Chemolithoautotrophic oxidation of thiosulfate, tetrathionate and thiocyanate by a novel rhizobacterium belonging to the genus *Paracoccus*. FEMS Microbiol. Lett. 270, 124–131. https://doi.org/10.1111/j.1574-6968.2007.00670.x.

Ghosh, W., Dam, B., 2009. Biochemistry and molecular biology of lithotrophic sulfur oxidation by taxonomically and ecologically diverse bacteria and archaea. FEMS Microbiol. Rev. 33, 999–1043. https://doi.org/10.1111/j.1574-6976.2009.00187.x.

Gomori, G., 1955. Preparation of buffers for use in enzyme studies. Methods Enzymol. 1, 138–146. https://doi.org/10.1016/0076-6879(55)01020-3.

Grimm, F., Franz, B., Dahl, C., 2008. Thiosulfate and Sulfur Oxidation in Purple Sulfur Bacteria, in: Dahl, C., Friedrich, C.G. (Eds.), Microbial Sulfur Metabolism. Springer, Berlin, Heidelberg, pp. 101–116. https://doi.org/10.1007/978-3-540-72682-1_9.

Hensen, D., Sperling, D., Trüper, H.G., Brune, D.C., Dahl, C., 2006. Thiosulphate oxidation in the phototrophic sulphur bacterium *Allochromatium vinosum*. Mol. Microbiol. 62, 794–810. https://doi.org/10.1111/j.1365-2958.2006.05408.x.

Houghton, J.L., Foustoukos, D.I., Flynn, T.M., Vetriani, C., Bradley, A.S., Fike, D.A., 2016. Thiosulfate oxidation by *Thiomicrospira thermophila*: metabolic flexibility in response to ambient geochemistry. Environ. Microbiol. 18, 3057–3072. https://doi.org/10.1111/1462-2920.13232.

Jørgensen, B.B., 1990a. A thiosulfate shunt in the sulfur cycle of marine sediments. Science, 249, 152–154. https://doi.org/10.1126/science.249.4965.152.

Jørgensen, B.B., 1990b. The sulfur cycle of freshwater sediments: role of thiosulfate. Limnol. Oceanogr. 35, 13291342. https://doi.org/10.4319/lo.1990.35.6.1329.

Kanao, T., Kamimura, K., Sugio, T., 2007. Identification of a gene encoding a tetrathionate hydrolase in *Acidithiobacillus ferrooxidans*. J. Biotechnol. 132, 16–22. https://doi.org/10.1016/j.jbiotec.2007.08.030.

Kanao, T., Kosaka, M., Yoshida, K., Nakayama, H., Tamada, T., Kuroki, R., Yamada, H., Takada, J., Kamimura, K., 2013. Crystallization and preliminary X-ray diffraction analysis of tetrathionate hydrolase from *Acidithiobacillus ferrooxidans*. Acta Cryst. F 69, 692–694. https://doi.org/10.1107/S1744309113013419.

Kappler, U., Friedrich, C.G., Trüper, H.G., Dahl, C., 2001. Evidence for two pathways of thiosulfate oxidation in *Starkeya novella* (formerly *Thiobacillus novellus*). Arch. Microbiol. 175, 102–111.https://doi.org/10.1007/s002030000241.

Kappler, U., Aguey-Zinsou, K.F., Hanson, G.R., Bernhardt, P.V., McEwan, A.G., 2004. Cytochrome *c*551 from *Starkeya novella* characterization, spectroscopic properties, and phylogeny of a diheme protein of the SoxAX family. J. Biol. Chem. 279, 6252–6260. https://doi.org/10.1074/jbc.M310644200.

Katayama, Y., Yasuhiko, M., Miyuki, K., Mai, K., Tadayoshi, M., Hiroshi, N., 1998. Cloning of genes coding for the three subunits of thiocyanate hydrolase of *Thiobacillus thioparus* THI 115 and their evolutionary relationships to nitrile hydratase. J. Bacteriol. 180, 2583–2589.

Kelly, D.P., Wood, A.P., 1994. Synthesis and determination of thiosulfate and polythionates. Methods Enzymol. 243, 475–501. https://doi.org/10.1016/0076-6879(94)43037-3.

Kelly, D.P., Shergill, J.K., Lu, W.P., Wood, A.P., 1997. Oxidative metabolism of inorganic sulfur compounds by bacteria. Antonie Van Leeuwenhoek. 71, 95–107. https://doi.org/10.1023/A:1000135707181.

Keltjens, J.T., Pol, A., Reimann, J., Op den Camp, H.J., 2014. PQQ-dependent methanol dehydrogenases: rare-earth elements make a difference. Appl. Microbiol. Biotechnol. 98, 6163–6183. https://doi.org/10.1007/s00253-014-5766-8.

Kikumoto, M., Nogami, S., Kanao, T., Takada, J., Kamimura, K., 2013. Tetrathionate-forming thiosulfate dehydrogenase from the acidophilic, chemolithoautotrophic bacterium *Acidithiobacillus ferrooxidans*. Appl. Environ. Microbiol. 79, 113–120. https://doi.org/10.1128/AEM.02251-12.

Kondo, R., Kasashima, N., Matsuda, H., Hata, Y., 2000. Determination of thiosulfate in a meromictic lake. Fish. Sci. 66, 1076–1081. https://doi.org/10.1046/j.1444-2906.2000.00171.x.

Kurth, J.M., Brito, J.A., Reuter, J., Flegler, A., Koch, T., Franke, T., Klein, E.M., Rowe, S.F., Butt, J.N., Denkmann, K., Pereira, I.A., Archer, M., Dahl, C., 2016. Electron accepting units of the diheme cytochrome *c* TsdA, a bifunctional thiosulfate dehydrogenase/tetrathionate reductase. J. Biol. Chem. 291, 24804–24818. https://doi.org/10.1074/jbc.M116.753863.

Kurtz, S., Phillippy, A., Delcher, A.L., Smoot, M., Shumway, M., Antonescu, C., Salzberg, S.L., 2004. Versatile and open software for comparing large genomes. Genome Biol, 5, R12. https://doi.org/10.1186/gb-2004-5-2-r12.

Luther, G.W., Giblin, A.E., Varsolona, R., 1985. Polarographic analysis of sulfur species in marine porewaters. Limnol. and Oceanogr. 30, 727–736.

Meulenberg, R., Pronk, J.T., Hazeu, W., van Dijken, J.P., Frank, J., Bos, P., Kuenen, J.G., 1993. Purification and partial characterization of thiosulphate dehydrogenase from *Thiobacillus acidophilus*. Microbiology 139, 2033–2039. https://doi.org/10.1099/00221287-139-9-2033.

Mopper, K., Kieber, D.J., 2002. Photochemistry and the cycling of carbon, sulfur, nitrogen and phosphorus, in: Dennis Hansell, D.A., Carlson, C.A., (Eds.), Biogeochemistry of marine dissolved organic matter, Academic Press, San Diego, CA, pp. 455–507. https://doi.org/10.1016/b978-012323841-2/50011-7.

Müller, F.H., Bandeiras, T.M., Urich, T., Teixeira, M., Gomes, C.M., Kletzin, A., 2004. Coupling of the pathway of sulphur oxidation to dioxygen reduction: characterization of a novel membrane-bound thiosulphate: quinone oxidoreductase. Mol. Microbiol. 53, 1147–1160. https://doi.org/10.1111/j.1365-2958.2004.04193.x.

Mukhopadhyaya, P.N., Deb, C., Lahiri, C., Roy, P., 2000. A *soxA* gene, encoding a diheme cytochrome *c*, and a *sox* locus, essential for sulfur oxidation in a new sulfur lithotrophic bacterium. J. Bacteriol. 182, 4278–4287. https://doi.org/10.1128/JB.182.15.4278-4287.2000.

Nakagawa, T., Mitsui, R., Tani, A., Sasa, K., Tashiro, S., Iwama, T., Hayakawa, T., Kawai, K., 2012. A catalytic role of XoxF1 as La^3^+-dependent methanol dehydrogenase in *Methylobacterium extorquens* strain AM1. PLoS One 7, e50480. https://doi.org/10.1371/journal.pone.0050480.

Orlova, M.V., Tarlachkov, S.V., Dubinina, G.A., Belousova, E.V., Tutukina, M.N., Grabovich, M.Y., 2016. Genomic insights into metabolic versatility of a lithotrophic sulfur-oxidizing diazotrophic Alphaproteobacterium *Azospirillum thiophilum*. FEMS Microbiol. Ecol. 92, fiw199. https://doi.org/10.1093/femsec/fiw199.

Owens, J.D., Reinhard, C.T., Rohrssen, M., Love, G.D., Lyons, T.W., 2016. Empirical links between trace metal cycling and marine microbial ecology during a large perturbation to Earth’s carbon cycle. Earth Planet. Sci. Lett. 449, 407–417. https://doi.org/10.1016/j.epsl.2016.05.046.

Parks, D.H., Imelfort, M., Skennerton, C.T., Hugenholtz, P., Tyson, G.W., 2015. CheckM: assessing the quality of microbial genomes recovered from isolates, single cells, and metagenomes. Genome Res. 25, 1043–1055. https://doi.org/10.1101/gr.186072.114.

Pol, A., Barends, T.R., Dietl, A., Khadem, A.F., Eygensteyn, J., Jetten, M.S., Op den Camp, H.J., 2014. Rare earth metals are essential for methanotrophic life in volcanic mudpots. Environ. Microbiol. 16, 255–264. https://doi.org/10.1111/1462-2920.12249.

Pyne, P., Alam, M., Ghosh, W., 2017. A novel soxO gene, encoding a glutathione disulfide reductase, is essential for tetrathionate oxidation in Advenella kashmirensis. Microbiol. res. 205, 1–7. https://doi.org/10.1016/j.micres.2017.08.002.

Pyne, P., Alam, M., Rameez, M.J., Mandal, S., Sar, A., Mondal, N., Debnath, U., Mathew, B., Misra, A.K., Mandal, A.K., Ghosh, W., 2018. Homologs from sulfur oxidation (Sox) and methanol dehydrogenation (Xox) enzyme systems collaborate to give rise to a novel pathway of chemolithotrophic tetrathionate oxidation. Mol. Microbiol. 109, 169–191. https://doi.org/10.1111/mmi.13972.

Quentmeier, A., Friedrich, C.G., 2001. The cysteine residue of the SoxY protein as the active site of protein-bound sulfur oxidation of *Paracoccus pantotrophus* GB17. FEBS Lett. 503, 168–172. https://doi.org/10.1016/S0014-5793(01)02727-2.

Rzhepishevska, O.I., Valdés, J., Marcinkeviciene, L., Gallardo, C.A., Meskys, R., Bonnefoy, V., Holmes, D.S., Dopson, M., 2007. Regulation of a novel *Acidithiobacillus caldus* gene cluster involved in metabolism of reduced inorganic sulfur compounds. Appl. Environ. Microbiol. 73, 7367–7372. https://doi.org/10.1128/AEM.01497-07.

Sambrook, J., Russell, D.W. 2001. Molecular Cloning: A Laboratory Manual. Cold Spring Harbor, N.Y Cold Spring Harbor Laboratory Press. p. A1.5.

Sauve, V., Bruno, S., Berks, B.C., Hemmings, A.M., 2007. The SoxYZ complex carries sulfur cycle intermediates on a peptide swinging arm. J. Biol. Chem. 282, 23194–204. https://doi.org/10.1074/jbc.M701602200.

Simon, R., Priefer, U., Pühler, A., 1983. A Broad Host Range Mobilization System for In Vivo Genetic Engineering: Transposon Mutagenesis in Gram Negative Bacteria. Nat. Biotechnol. 1, 784–791. https://doi.org/10.1038/nbt1183-784.

Skorupski, K., Taylor, R.K., 1996. Positive selection vectors for allelic exchange. Gene, 169, 47–52. https://doi.org/10.1016/0378-1119(95)00793-8.

Skovran, E., Martinez-Gomez, N.C., 2015. Just add lanthanides. Science 348, 862–863. https://doi.org/10.1126/science.aaa9091.

Sorokin, D. Y., Tourova, T. P., Lysenko, A. M., Kuenen, J. G., 2001. Microbial thiocyanate utilization under highly alkaline conditions. Appl. Environ. Microbiol. 67, 528–538. https://doi.org/10.1128/AEM.67.2.528-538.2001.

Sung, Y.C., Fuchs, J.A., 1988. Characterization of the *cyn* operon in *Escherichia coli* K12. J. Biol. Chem. 263, 14769–14775.

Thamdrup, B., Finster, K., Fossing, H., Hansen, J.W., Jørgensen, B.B., 1994. Thiosulfate and sulfite distributions in porewater of marine sediments related to manganese, iron, and sulfur geochemistry. Geochim. Cosmochim. Acta 58, 67–73. https://doi.org/10.1016/0016-7037(94)90446-4.

Tsallagov, S.I., Sorokin, D.Y., Tikhonova, T.V., Popov, V.O., Muyzer, G., 2019. Comparative Genomics of *Thiohalobacter thiocyanaticus* HRh1T and *Guyparkeria* sp. SCN-R1, Halophilic Chemolithoautotrophic Sulfur-Oxidizing *Gammaproteobacteria* Capable of Using Thiocyanate as Energy Source. Front. Microbiol.10, Article 898. https://doi.org/10.3389/fmicb.2019.00898.

van Zyl, L.J., van Munster, J.M., Rawlings, D.E., 2008. Construction of *arsB* and *tetH* mutants of the sulfur-oxidizing bacterium *Acidithiobacillus caldus* by marker exchange. Appl. Environ. Microbiol. 74, 5686–5694. https://doi.org/10.1128/AEM.01235-08.

Verte, F., Kostanjevecki, V., De Smet, L., Meyer, T., Cusanovich, M., Van Beeumen, J., 2002. Identification of a thiosulfate utilization gene cluster from the green phototrophic bacterium *Chlorobium limicola*. Biochemistry 41, 2932–2945. https://doi.org/10.1021/bi011404m.

Visser, J.M., de Jong, G.A., Robertson, L.A., Kuenen, J.G., 1996. Purification and characterization of a periplasmic thiosulfate dehydrogenase from the obligately autotrophic *Thiobacillus* sp. W5. Arch. Microbiol.166, 372-378. https://doi.org/10.1007/BF01682982.

West, P.W., Gaeke, G.C., 1956. Fixation of sulfur dioxide as disulfitomercurate (II) and subsequent colorimetric estimation. Anal. Chem. 28, 1816–1819. https://doi.org/10.1021/ac60120a005.

Xu, Y., Schoonen, M.A.A., Nordstrom, D.K., Cunningham, K.M., Ball, J.W., 1998. Sulfur geochemistry of hydrothermal waters in Yellowstone National Park: I The origin of thiosulfate in hot spring waters. Geochim Cosmochim Acta. 62, 3729–3743. https://doi.org/10.1016/S0016-7037(98)00269-5.

