## Supplemental information for "Two pathways for thiosulfate oxidation in the alphaproteobacterial chemolithotroph *Paracoccus thiocyanatus* SST"

**Running title:**	Thiosulfate oxidation via tetrathionate in an alphaproteobacterium

**Keywords:**	sulfur**-**chemolithotrophy, Sox multienzyme system, Alphaproteobacteria, thiosulfate oxidation via tetrathionate-intermediate, thiosulfate dehydrogenase, tetrathionate oxidation

**List of the materials included:**

1. Supporting tables:
   i) Tables S1 through S4, with relevant titles and notes.
2. Supporting figures:
   i) Figures S1 through S4, with respective figure legends.
3. Supporting references.

**Supplementary Tables**

**Table S1. Primers used in this study for the knock-out mutagenesis of tsdA.** The *Xba*I site is in bold font, the *EcoR*I site is in bold underlined font, while *Kpn*I site is underlined.

| Primers | Sequence (5՛-3՛) |
| --- | --- |
| Pt _Ko_tsdA_F | GTTT**GAATTC**AATTGCTCGACCTGCCACC |
| Pt _Ko_tsdA_R | TTT**TCTAGA**ACAGCACCACCCCATCCATGC |
| Pt_Ko_tsdA_FU_F | GCGAAGATGAGCTCTGAACATCACGACGATACGGTACCTATTACGGGCAACAGGTGAACG |
| Pt_Ko_tsdA_FU_R | ATCGTCGTGATGTTCAGAGCTCATCTTCGC |
| Pt_Ko_tsdA_F2 | CGACATCTTCATGAACACCTCGACCAATGC |
| Pt_Ko_tsdA_F3 | CGCAGCAGCCCGCCTATGTCGCCAGC |
| Pt _Ko_tsdA_R2 | AACCATCGACCACATGCCGGCAAAGACG |
| kanR_KpnI_F | TAGACGGTACCGTTTTATGGACAGCAAGCG |
| kanR_KpnI_R | GGGGTACCAAGAACTCCAGCATGAGATCCC |

**Table S2.Primers used in is study for the knock-out mutagenesis of soxB.** The XbaI site is in bold font, the SacI site is in bold underlined font, while KpnI site is underlined.

| Primers | Sequence (5՛-3՛) |
| --- | --- |
| Pt_Ko_soxB_F | TAAGC**GAGCTC**CTACAAGGACCAGATCCAGG |
| Pt_Ko_soxB_R | TTT**TCTAGA**GGAAGGAACGACCGTTGGACG |
| Pt_Ko_soxB_FU_F | CCAGTTGAAGCCGATCTATTTCCGCGAACCATCAGGTACCAGGACATCCACAACGTGACC |
| Pt_Ko_soxB_FU_R | GGTTCGCGGAAATAGATCGGCTTCAACTGG |
| Pt_Ko_soxB_R2 | ATTTCCTTGACCCGGTCGGTGCCATAGG |
| Pt _Ko_soxB _F2 | CGCACAGGAGTGACCGAGGAATGATTACCC |
| Pt _Ko_soxB _R3 | GCCGTCCTTGGTGAGTTCGTCTTTCATGG |
| kanR_KpnI_F | TAGACGGTACCGTTTTATGGACAGCAAGCG |
| kanR_KpnI_R | GGGGTACCAAGAACTCCAGCATGAGATCCC |

**Table S3. Primers used in this study for the knock-out mutagenesis of xoxF.** The XbaI site is in bold font, the SacI site is in bold underlined font, while KpnI site is underlined.

| Primers | Sequence (5՛-3՛) |
| --- | --- |
| Pt_Ko_xoxF_F | CCG**TCTAGA**GTGCTTGCGCAGGAAGACACG |
| Pt_Ko_xoxF_R | TTT**GAGCTC**ACCAGCAATTCACCGGTTTCG |
| Pt _Ko_xoxF_FU_F | GGTGA**GGTACC**CCAACAAGATCGGCGATCC |
| Pt _Ko_xoxF_FU_R | GGATCGCCGATCTTGTTGGGGTACCTCACC |
| kanR_KpnI_F | TAGACGGTACCGTTTTATGGACAGCAAGCG |
| kanR_KpnI_R | GGGGTACCAAGAACTCCAGCATGAGATCCC |

**Table S4. Optimum pH for organotrophic growth of *P. thiocyanatus* SST in Luria Bertani medium.** Luria Bertani (LB) medium was inoculated with 1% (v/v) LB-culture of *P. thiocyanatus* SST having OD_600_ of 0.6. Growth in terms of biomass production was compared in the form of absorbance at OD_600_.

| Incubation time (Hours) |  | pH5 | pH 6 | pH 7 | pH 7.5 | pH 8 | pH 8.5 | pH 9 | pH 10 | pH 11 | B |
| --- | --- | --- | --- | --- | --- | --- | --- | --- | --- | --- | --- |
| 12 | OD | 0 | 0.0238 | 0.4036 | 0.4397 | 0.4493 | 0.4349 | 0.3191 | 0.005 | 0.0045 | 0 |
| 24 | OD | 0 | 0.1319 | 1.7929 | 1.8368 | 1.8629 | 1.8032 | 1.4087 | 0.035 | - | - |

**Supplementary Figures**

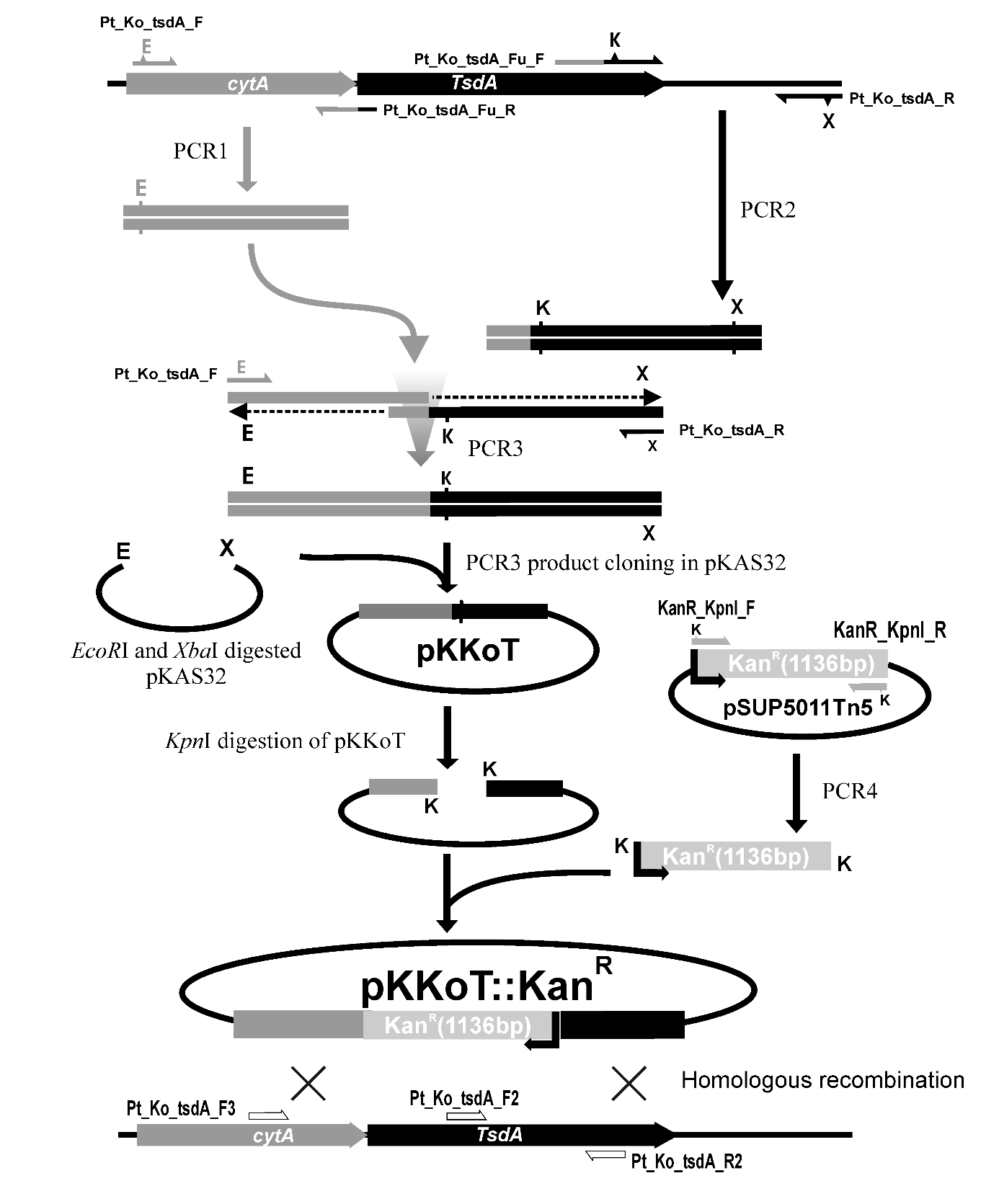

**Fig. S1. Schematic representation of steps involving knockout mutagenesis of tsdA homolog of SST via replacement with Kan^R^ cartridge.** Restriction sites XbaI, EcoRI and KpnI are designated as X, E and K.Homologous regions to be involved in recombination events are marked by cross mark between them. Primers used for mutant confirmation by PCR and sequencing are colored white and shownin the bottom of the figure at tentative position beside the genomic locus to be mutated.

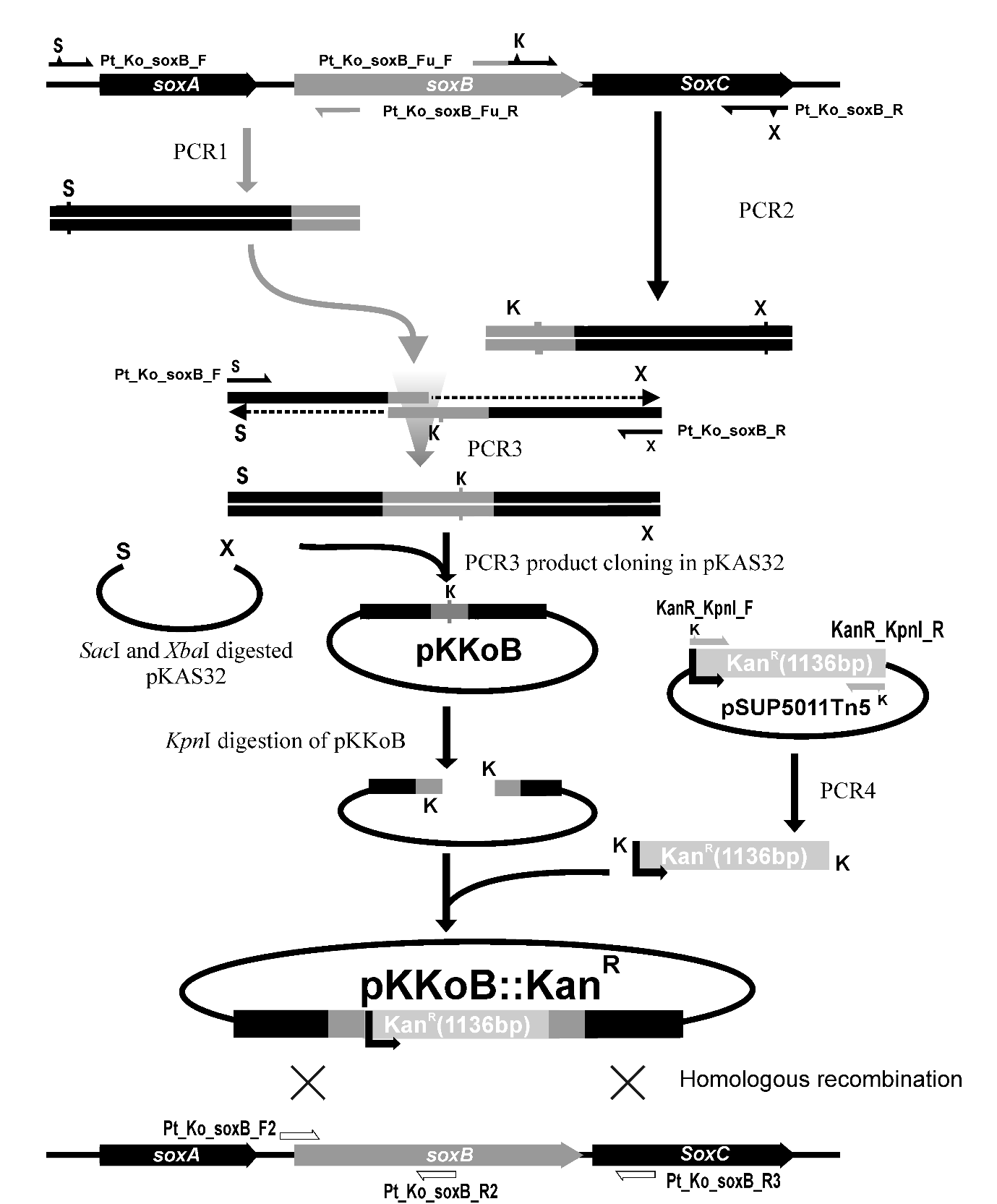

**Fig. S2. Schematic representation of steps involving knockout mutagenesis of soxB homologof SST via replacement with KanRcartridge.** Restriction sites *Xba*I, *Sac*I and *Kpn*I are designated as X, S and K.Homologous regions to be involved in recombination events are marked by cross mark between them. Primers used for mutant confirmation by PCR and sequencing are colored white and shown in the bottom of the figure at tentative position beside the genomic locus to be mutated.

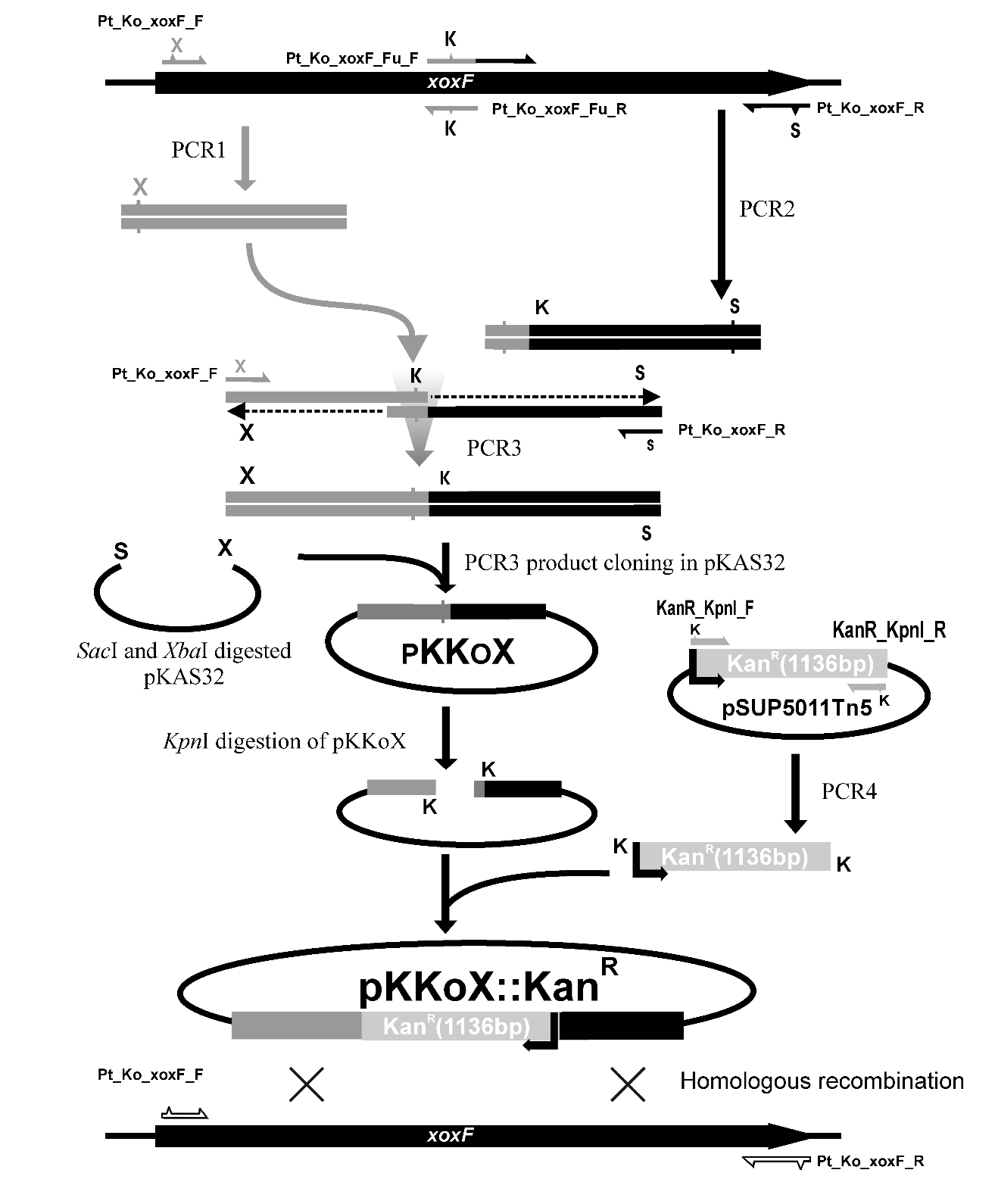

**Fig. S3. Schematic representation steps involving knockout mutagenesis of xoxF homologvia insertion ofKan^R^cartridge inside the ORF.** Restriction sites XbaI, SacI and KpnI are designated as X, S and K. Homologous regions to be involved in recombination events are marked by cross mark between them. Primers used for mutant confirmation by PCR and sequencing are colored white and shown in the bottom of the figure at tentative position beside the genomic locus to be mutated.

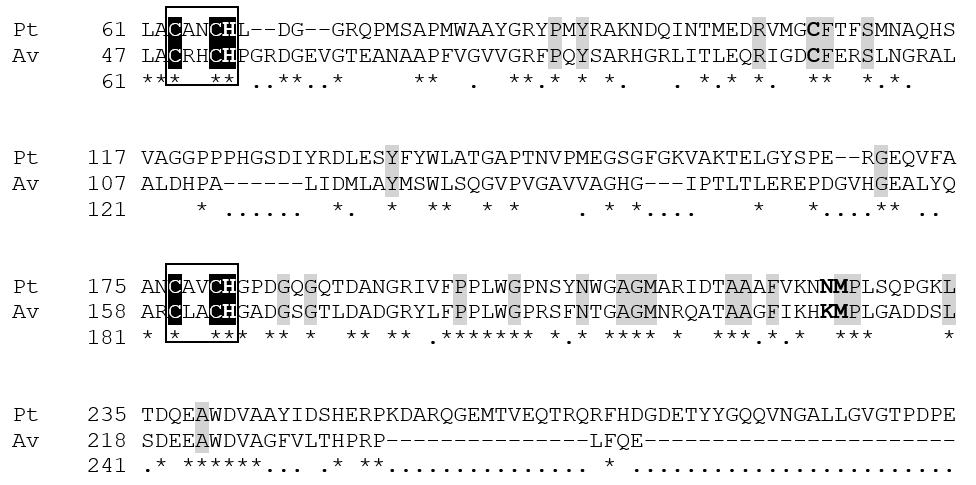

**Fig. S4. TsdA of P. thiocyanatus SST contains heme binding residues required for thiosulfate to tetrathionate conversion**. Alignment of the primary sequence of the TsdA homologs of P. thiocyanatus (WP_020423861.1), Pt and Allochromatium vinosum (WP_012969337.1), Av. The consensus CXXCH heme-attachment motifs (Denkmann et al., 2012) are enclosed in boxes. The conserved ligand residues axially co-ordinating the heme 1 (His^53^ and Cys^96^) and heme 2 (His^164^ and Lys^208^/Met^209^) are shown in bold (Brito et al., 2015). Residues conserved in homologs across all classes of proteobacteria (Denkmannet al., 2012) are highlighted in gray. Identical residues are marked with asterisks. The numberings on the individual sequences refer to the amino acid positions in their respective complete primary sequences after removing signal peptide sequences as predicted by SignalP 3.0 (http://www.cbs.dtu.dk/services/SignalP-3.0/).

Supplementary references

Brito, J.A., Denkmann, K., Pereira, I.A.C., Archer,M., Dahl, C., 2015. Thiosulfate Dehydrogenase (TsdA) from *A. vinosum*: structural and functional insights into thiosulfate oxidation. J. Biol. Chem. 290, 9222-9238. https://doi.org/10.1074/jbc.M114.623397.

Denkmann, K., Grein, F., Zigann,R., Siemen, A., Bergmann, J., van Helmont, S., Nicolai, A., Pereira, I.A.C., Dahl, C., 2012. Thiosulfate dehydrogenase: a widespread unusual acidophilic c‐type cytochrome. Environ. Microbiol. 14, 2673-2688. https://doi.org/10.1111/j.1462-2920.2012.02820.x.

Nielsen, H., Engelbrecht, J., Brunak, S., Heijne, G.V., 1997. Identification of prokaryotic and eukaryotic signal peptides and prediction of their cleavage sites. Protein Eng. Des. Sel. 10, 1-6.<https://doi.org/>

10.1093/protein/10.1.1.
